# The CP110-CEP97-CEP290 module orchestrates a centriolar satellite dependent response to proteotoxic stress

**DOI:** 10.1101/2020.12.20.423672

**Authors:** Suzanna L. Prosser, Johnny Tkach, Ladan Gheiratmand, Ciaran G. Morrison, Laurence Pelletier

## Abstract

Protein degradation at the centrosome, the primary microtubule organizing centre of the cell, is critical to a myriad of cellular processes. Perturbation of the ubiquitin proteasome system causes the formation of an inclusion, or aggresome, at the centrosome. By systematic microscopy analysis, we have placed a subset of centrosomal proteins within the aggresome. Centriolar satellites, proteinaceous granules found in the vicinity of centrosomes, also became incorporated into this structure. Through high-resolution quantitative analysis, we have defined aggresome assembly at the centrosome, demonstrating a requirement for satellites in this process. Furthermore, a module consisting of CP110-CEP97-CEP290 was required to recruit aggresome components early in the pathway and senescent cells were defective in aggresome formation due to limiting amounts of CP110. Finally, satellites and the CP110-CEP97-CEP290 module were required for the aggregation of mutant huntingtin. The accumulation of protein aggregates is central to the pathology of a range of human disorders. These data thereby reveal new roles for CP110, its interactors, and centriolar satellites in controlling cellular proteostasis and the aggregation of disease relevant proteins.

## INTRODUCTION

Centrosomes serve as both the main microtubule organizing centre and template for primary cilia formation, thereby involving them in multiple critical cellular functions (Conduit et al., 2015; Joukov and De Nicolo, 2019). The centrosome is a highly dynamic non-membrane bound organelle consisting of a pair of centrioles surrounded by pericentriolar material (PCM). In addition, a number of proteinaceous granules, called centriolar satellites, are found in close proximity to the centrosome (Kubo et al., 1999). Satellites are required for the correct assembly and functioning of centrosomes, and are implicated in multiple processes throughout the cell (Gheiratmand et al., 2019; Odabasi et al., 2019; Quarantotti et al., 2019). The centrosome contributes to a range of diverse cellular events in part through acting as a scaffold for localized protein degradation via the Ubiquitin Proteasome System (UPS; Kimura et al., 2014; Vora and Phillips, 2016). UPS machinery complexes localize to the centrosome (Fabunmi et al., 2000; Wigley et al., 1999), with protein degradation at the centrosome regulating processes such as cell cycle control (Clute and Pines, 1999; Máthé et al., 2004; Raff et al., 2002), cell fate acquisition (Fuentealba et al., 2008; 2007; Vora and Phillips, 2015), neuronal cell morphogenesis (Kim et al., 2009; Puram et al., 2010; 2013), immune system function (Antón et al., 1999; Hung et al., 2003; Lacaille and Androlewicz, 2000), and cellular stress responses (Vertii et al., 2015; Vidair et al., 1993). However, the specific function of centrosomal components in the regulation of proteostasis remains unclear.

If the capacity of the UPS is exceeded, through a decrease in proteasomal activity or increase in misfolded polypeptides, cells accumulate proteins into an inclusion, called the aggresome, at the centrosome (Johnston et al., 1998). The organization of potentially toxic protein species into a single location serves a protective function, while also facilitating their clearance by autophagy (Fortun et al., 2003; Hao et al., 2013; Wong et al., 2008). Aggresome formation is an active process that requires intact microtubules, the motor protein dynein, and the microtubule deacetylase HDAC6 (García-Mata et al., 1999; Johnston et al., 2002; Kawaguchi et al., 2003). In addition to ubiquitinated proteins, aggresomes are rich in proteasomal subunits, chaperone proteins, ubiquitin ligases and ubiquitin-binding proteins (Choi et al., 2020; Fusco et al., 2012; García-Mata et al., 1999; Mao et al., 2017; Mishra et al., 2009; Wigley et al., 1999; Zhang and Qian, 2011; Zhou et al., 2014b). Some centrosomal proteins have also been shown to accumulate in aggresomes (Chiba et al., 2012; Johnston et al., 1998; McNaught et al., 2002; Szebenyi et al., 2007), along with PCM1, the major structural component of centriolar satellites (Didier et al., 2008). Similar to the aggresomal assembly pathway, satellites require microtubules and dynein for their association with centrosomes (Dammermann and Merdes, 2002; Kubo et al., 1999). Growing evidence implicates satellites as mediators of protein stability via both autophagy and the UPS (Joachim et al., 2017; Wang et al., 2016). The satellite proteins OFD1 and BBS4 interact with UPS subunits, with depletion of either leading to the loss of proteasomal machinery from the centrosome and deficiencies in proteasome-dependent pathways (Gerdes et al., 2007; Liu et al., 2014). Moreover, changes in the cellular proteome in satellite-deficient cells highlights satellites’ function as regulators of global proteostasis (Odabasi et al., 2019). However, although aggresome assembly occurs at the centrosome, the role of centriolar satellites in aggresome formation has yet to be explored.

Centrosome biology itself is tightly regulated by proteasomal degradation (Cunha-Ferreira et al., 2009; Fung et al., 2018; Guderian et al., 2010; Hames et al., 2005; Nigg and Holland, 2018; Puklowski et al., 2011). Strict control over the abundance of centriole duplication factors, such as PLK4, SAS6 and STIL, stringently regulates centriole number (Arquint et al., 2012; Korzeniewski et al., 2009; Rogers et al., 2009; Strnad et al., 2007), with centriole amplification observed following prolonged inhibition of the proteasome (Duensing et al., 2007). In human cells, centriole overduplication induced by PLK4 overexpression requires the centriolar distal end protein CP110 (Kleylein-Sohn et al., 2007). Conversely, in flies, normal levels of CP110 in primary spermatocytes suppresses centriole overduplication induced by overexpression of the *Drosophila* homolog of PLK4 (Franz et al., 2013). Proteasomal inhibition also leads to the appearance of elongated centrioles (Korzeniewski et al., 2010). Centriole length is regulated by a number of proteins, including CP110 (Schmidt et al., 2009). CP110 and its partner CEP97 form a cap at the centriole distal end that restricts centriole length, with depletion of either protein resulting in elongated centriole structures (Kohlmaier et al., 2009; Schmidt et al., 2009; Spektor et al., 2007). CP110 and CEP97 also act as a block to primary cilium formation, as their removal from the mother centriole is a prerequisite for axonemal microtubule extension during ciliogenesis (Čajánek and Nigg, 2014; Goetz et al., 2012; Tsang et al., 2008). This ciliation suppressor function of CP110 is dependent upon its interaction with the satellite protein CEP290 (Tsang et al., 2008). However, CP110 can promote ciliogenesis, as CP110-deficient mice exhibit ciliary deficiencies (Yadav et al., 2016). Levels of CP110 are tightly regulated via the antagonistic actions of the SCF^CyclinF^ ubiquitin ligase complex (D’Angiolella et al., 2010), and the deubiquitinating enzyme (DUB) USP33 (Li et al., 2013), suggesting proteolytic regulation is vital to CP110’s role(s) in the cell. Of note, CP110 and CEP97 were recently reported in the centriolar satellite proteome and interactome (Gheiratmand et al., 2019; Quarantotti et al., 2019). Whether the function of CP110 and its partners extends beyond the control of centriole number and elongation is unknown.

Senescence describes the process by which cells cease dividing and undergo distinctive phenotypic changes (van Deursen, 2014). The number of senescent cells in an organism increases with age, with senescence contributing to age-related pathologies such as cancer, tissue degeneration, inflammatory disease and neurodegeneration. Centrosome dysfunction is believed to contribute to the senescence process (Wu et al., 2020). Proteasomal activity decreases during senescence, resulting in the accumulation of damaged and misfolded proteins that impair cellular functions, features that are associated with late-onset pathologies (Cuanalo-Contreras et al., 2013; Fernández-Cruz and Reynaud, 2020; López-Otín et al., 2013; Saez and Vilchez, 2014). The accumulation of misfolded proteins into proteinaceous inclusions is a hallmark of neurodegenerative disorders, including Huntington’s disease (HD). The presence of expanded polyglutamine (polyQ) tracts in the HD protein huntingtin (HTT) are associated with protein aggregation and inclusion formation (Adegbuyiro et al., 2017). HTT interacts with PCM1 through huntingtin-associated protein 1 (HAP1) and expression of polyQ-HTT causes PCM1 to aggregate at the centrosome (Keryer et al., 2011). The potential roles of satellites in the aggregation of polyQ-HTT remain to be determined.

Here, we describe the association of a set of centriolar and PCM proteins with the aggresome. Centriolar satellites also accumulated into the aggresome in response to proteasomal inhibition and aggresome assembly was reduced in cells with disrupted satellites. Furthermore, a module consisting of CP110, CEP97 and CEP290 was found to operate early in the aggresome assembly pathway. Finally, we provide evidence to show that aggresome formation is defective in senescent cells due to limiting levels of CP110, and implicate satellites and the CP110-CEP97-CEP290 module in the formation of polyQ-HTT inclusions.

## RESULTS

### Centrosome and centriolar satellite proteins localize to the aggresome upon proteasome inhibition

Short-term treatment of RPE-1 cells with the proteasome inhibitors MG132 and bortezomib (BZ) led to the accumulation of ubiquitinated proteins (Ub^+^) at the centrosome (Fig. 1A-B). Accumulation of proteins that are known to incorporate into the aggresome, such as pHSP27, HDAC6, p62, HSP70, HAP1, and dynein, confirmed that we were inducing aggresome assembly (Fig. S1A; Bolhuis and Richter-Landsberg, 2010; Fujinaga et al., 2009; García-Mata et al., 1999; Johnston et al., 2002; Zhou et al., 2014b). Over 75% of cells treated with proteasome inhibitors for 5 hours responded by forming an aggresome (Fig. 1C). Furthermore, aggresome formation was universal in additional cell lines (A-375, BJ-5ta, HFF-1, HeLa, and U-2 OS) tested in this study (Fig. S1B-C).

**Figure 1.**
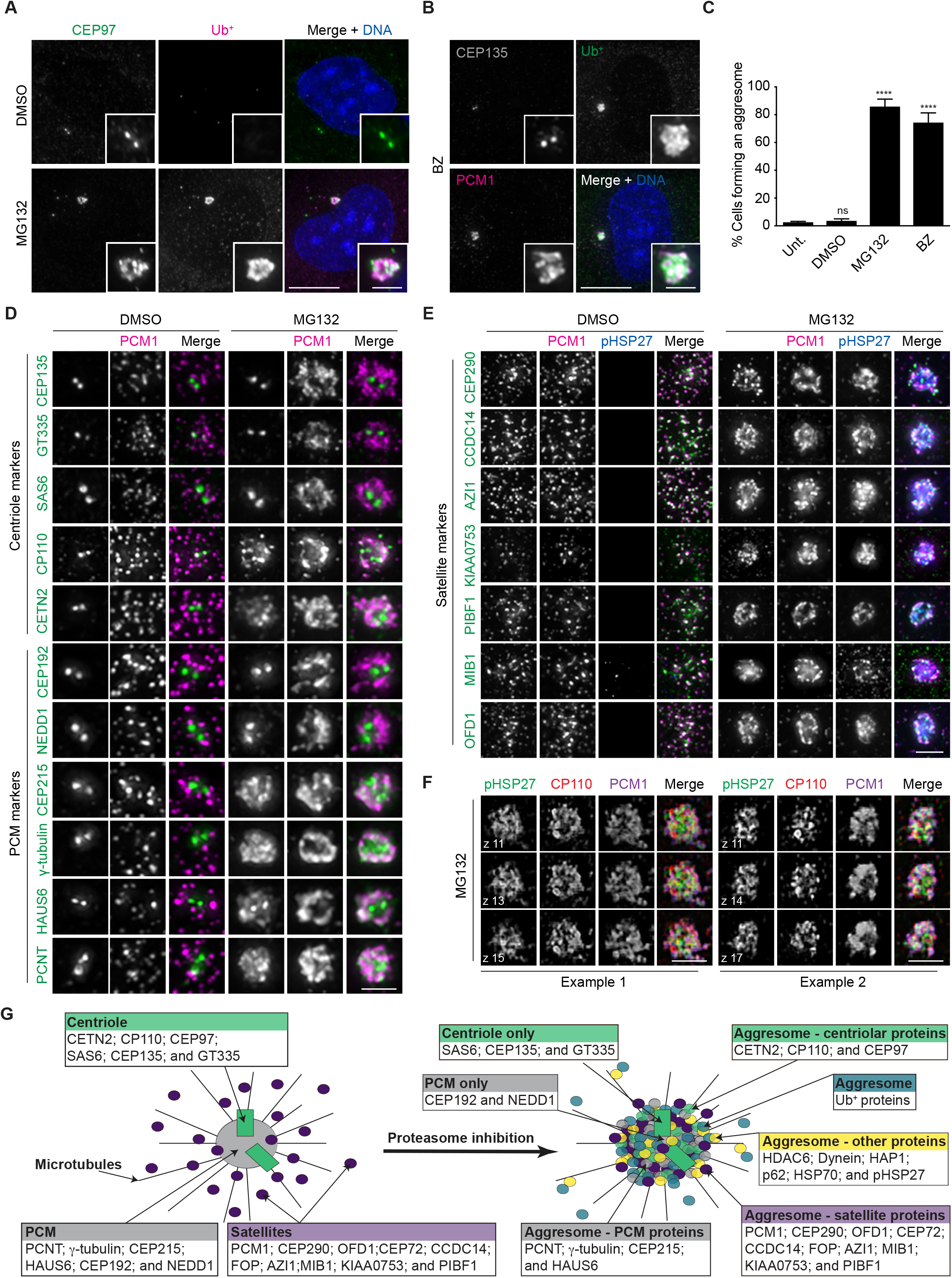
Centrosomal proteins localize to the aggresome upon proteasome inhibition. **A**. RPE-1 cells treated with DMSO or MG132 were stained for CEP97, ubiquitinated (Ub^+^) proteins and DNA (DAPI). **B**. RPE-1 cells treated with bortezomib (BZ) were for CEP135, Ub^+^ proteins, PCM1 and DNA (DAPI). **C**. Histogram showing the percentage of cells forming an aggresome in untreated (Unt.), DMSO, MG132 and BZ treated cells as revealed by Ub^+^ staining. **D**. RPE-1 cells treated with DMSO or MG132 were stained with antibodies against PCM1 and the indicated protein. **E**. RPE-1 cells treated with DMSO or MG132 were stained with antibodies against PCM1, pHSP27 and the indicated protein. **F**. Higher resolution images of aggresomes in MG132 treated RPE-1 cells stained for pHSP27, CP110 and PCM1. Three individual z-planes are shown for each example. **G**. Schematic showing proteins localizing to the centrosome and their redistribution to the aggresome upon proteasome inhibition. Scale bars: A and B 10 μm, insets 2 μm; D, E and F 2 μm. ****p < 0.0001; ns, not significant. See also Figure S1.

Given the recruitment of CEP97 and PCM1 to the aggresome here (Fig. 1A and B) and a previous report that γ-tubulin, PCNT, ninein, and PCM1 accumulate at the centrosome during proteasome inhibition (Didier et al., 2008), we sought to systematically survey the localization of centrosomal proteins following MG132 treatment. To that end, we stained for centriole (CEP135, GT335, SAS6, CP110 and CETN2) and PCM (CEP192, NEDD1, CEP215, γ-tubulin, HAUS6 and PCNT) markers (Fig. 1D). Similar to CEP97, CP110 and CETN2 associated with the aggresome. However, CEP135, GT335 and SAS6 remained restricted to the centrioles. CEP215, γ-tubulin, HAUS6 and PCNT also localized to the aggresome, with CEP192 and NEDD1 staying limited to the PCM.

PCM1 is a major component of centriolar satellites (Kubo et al., 1999). We therefore examined whether other satellite proteins are also recruited to the aggresome. All of the satellite proteins we surveyed (CEP290, CCDC14, AZI1, KIAA0753, PIBF1, MIB1, OFD1, CEP72, and FOP) localized with pHSP27 and PCM1 in the aggresome (Fig. 1E and Fig. S1D). Further to this, higher resolution imaging revealed that pHSP27, CP110 and PCM1 occupy distinct, but overlapping, spatial domains within the aggresome (Fig. 1F and S1E). In summary, a subset of centriolar and PCM proteins localize to the aggresome following proteasome inhibition, along with centriolar satellites (Fig. 1G).

### High-resolution quantitative analysis reveals that centriolar satellites are required for aggresome formation

As centriolar satellites localize to the aggresome, we wondered whether satellites contribute to aggresome assembly. To examine the aggresome pathway, we established an automated image analysis pipeline to measure aggresome area and satellite distribution (Fig. S2A; see Methods). pHSP27 was chosen as the aggresome marker for this analysis due to its low background in control-treated cells and distinct signal around the centrosome in cells treated with MG132 (Fig. S2B). Knockdown of HSP27 demonstrated the specificity of this signal (Fig. S2B-C), with our analysis detecting a reduction of pHSP27 to control levels in cells depleted of HSP27 (Fig. S2D). Staining for Ub^+^ proteins and CP110 revealed that this led to a block to aggresome assembly, with Ub^+^ labelling appearing restricted to a ring around each centriole (Fig. S2E). To map satellite distribution in cells, we used PCM1 labelling, which revealed a distinct movement of satellites into the aggresome region following treatment with MG132, with 90.1% of satellites in close proximity to the centrosome as compared to 68.1% in controls (Fig. S2F). Furthermore, satellites still migrated into the pericentrosomal region in HSP27 siRNA-treated cells, despite aggresome formation being blocked (Fig. S2F).

To further validate our quantification protocol, we utilized approaches previously reported to disrupt aggresome formation, but until now had not been subjected to rigorous analysis. First, we used cycloheximide (CHX) to inhibit protein translation as active translation is required for aggresome assembly (Meriin et al., 2012; Nawrocki et al., 2005; Qin et al., 2019; Wójcik et al., 1996; Zhou et al., 2014a). A 30-minute pre-treatment with CHX before proteasome inhibition completely blocked pHSP27 and Ub^+^ recruitment to the aggresome (Fig. 2A-C). Strikingly, satellites also failed to accumulate in the pericentrosomal region in cells treated with CHX and MG132 (Fig. 2D).

**Figure 2.**
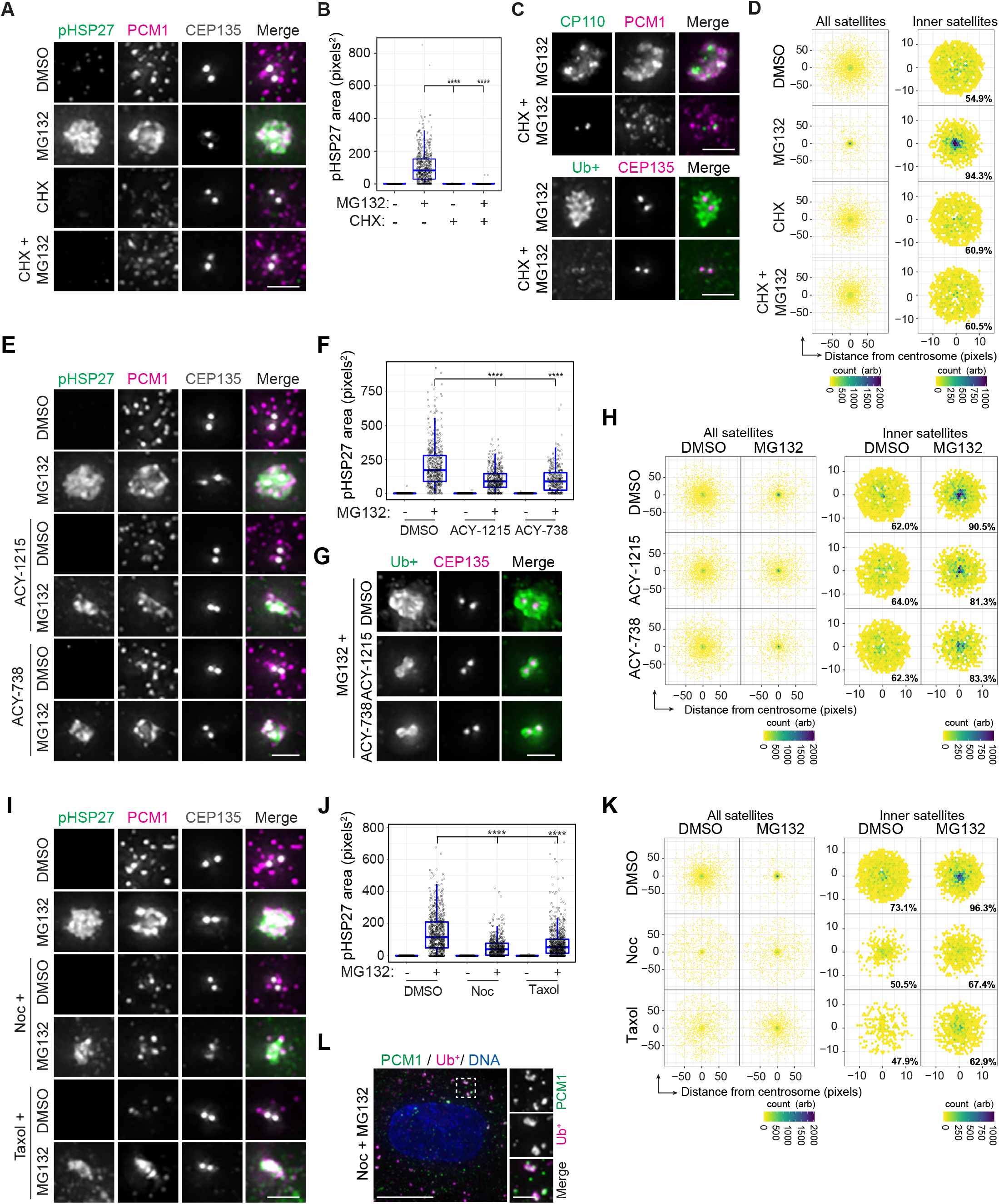
High-resolution quantitative analysis confirms the requirement for protein translation, HDAC6 and microtubules in aggresome formation. **A**. RPE-1 cells treated with DMSO, MG132 or cycloheximide (CHX) alone, or CHX for 30 minutes followed by MG132, were stained for pHSP27, PCM1 and CEP135. **B**. Box-and-whisker plot showing the area occupied by pHSP27 in cells treated as in A. **C**. Staining of PCM1 with CP110 or Ub^+^ proteins in cells treated with MG132 +/− pre-treatment with CHX. **D**. Intensity maps of PCM1 distribution relative to the centrosome in cells treated as in A. The percentage PCM1 signal residing in the defined ‘inner’ region is indicated. **E**. DMSO and MG132 treated RPE-1 cells were treated concurrently with ACY-1215 or ACY-738 as indicated, then stained for pHSP27, PCM1 and CEP135. **F**. Box-and-whisker plot showing the area occupied by pHSP27 in cells treated as in E. **G**. RPE-1 cells treated with MG132 +/− ACY-1215 or ACY-738 were stained for Ub^+^ proteins and CEP135. **H**. Intensity maps of PCM1 distribution relative to the centrosome in cells treated as in E. **I**. RPE-1 cells were pre-treated with nocodazole (Noc) or taxol for 2 hours before treatment with DMSO or MG132. Cells were fixed and stained for pHSP27, PCM1 and CEP135. **J**. Box-and-whisker plot showing the area occupied by pHSP27 in cells treated as in I. **K**. Intensity maps of PCM1 distribution relative to the centrosome in cells treated as in I. **L**. RPE-1 cells treated with MG132 and nocodazole were stained for PCM1, Ub^+^ proteins and DNA (DAPI). Scale bars: A, C, E, G and I 2 μm; L 10 μm, inset 2 μm. ****p < 0.0001. See also Figure S2.

Next, we used two pharmacological inhibitors of HDAC6, ACY-1215 and ACY-738 (Wang et al., 2018), as aggresome assembly requires HDAC6 activity (Kawaguchi et al., 2003). Inhibition was confirmed by an increase in the levels of acetylated tubulin (Fig. S2G). Quantification of aggresome formation in cells treated with either HDAC6 inhibitor and MG132 revealed a decrease in aggresome size, with pHSP27 restricted to the area immediately around the centrioles (Fig. 2E-F) and Ub^+^ proteins restricted to a ring around each centriole (Fig. 2G). However, the accumulation of satellites in response to MG132 was largely unaffected by HDAC6 inhibition (Fig. 2H).

Finally, we used nocodazole to depolymerize, and taxol to stabilize, microtubules as their dynamic instability is required for aggresome formation (Bauer and Richter-Landsberg, 2006). Treatment of cells with either compound led to the dispersal of satellites throughout the cytoplasm (Fig. S2H; Conkar et al., 2017; Dammermann and Merdes, 2002; Kubo et al., 1999). When cells were also treated with MG132, aggresomal pHSP27 accumulation was reduced and satellite recruitment restricted (Fig. 2I-K). Ub^+^ proteins in these cells formed multiple aggregates throughout the cytoplasm, which colocalized with PCM1 (Figs. 2L and S2I), suggesting satellites associate with Ub^+^ proteins in the cytoplasm before they are assembled into an aggresome. In cells that lack centrioles through the disruption of STIL, Ub^+^ proteins similarly colocalized with PCM1 foci in the cytoplasm (Fig. S2J), demonstrating that this association was not dependent upon a functional centrosome. However, cytoplasmic Ub^+^ aggregates in cells treated with ACY-1215 did not contain PCM1 (Fig. S2K), suggesting that HDAC6 activity is required for this association. Treatment of cells with nocodazole after an aggresome had formed revealed that microtubules are not required for the maintenance of this structure or the association of satellites with it after it has assembled (Fig. S2L-N). Together, these results demonstrate that our quantification pipeline can robustly measure aggresomes and is sensitive enough to distinguish between complete and partial blocks to aggresome assembly. Furthermore, satellites move towards the centrosome in response to proteasomal inhibition, downstream of the requirement for active protein translation in the aggresome pathway.

To test the requirement for satellites in aggresome assembly, we used CRISPR/Cas9 to disrupt the genes encoding the satellite proteins AZI1/CEP131, CCDC14, KIAA0753 and PIBF1/CEP90 in RPE-1 cells. Gene disruption was confirmed by genomic PCR and sequence analysis using ICE (Inference of CRISPR Edits, Synthego; Table S1) and protein loss demonstrated by immunoblot and immunofluorescence (IF) microscopy (Fig. S3A-B). In each knockout (KO) line, PCM1 positive-satellites persisted (Fig. S3B), while in a previously generated PCM1 KO line AZI1, CCDC14, KIAA0753 and PIBF1 were restricted to the centrosomes (Fig. S3B; Gheiratmand et al., 2019), indicative of satellite loss in these cells, and in line with previous studies (Hoang-Minh et al., 2016; Odabasi et al., 2019; Wang et al., 2016).

The ability to form an aggresome was reduced to a similar extent in each KO line, despite satellites accumulating in the pericentrosomal region of AZI1, CCDC14, KIAA0753 and PIBF1-depleted cells (Figs. 3A-C and S3C). As seen for other methods that block aggresome formation, Ub^+^ proteins were restricted to a ring around each centriole (Fig. 3D) and other aggresome markers, including HDAC6, p62, HSP70, HAP1 and CP110, failed to accumulate (Fig. 3E). The effect of satellite disruption on aggresome formation was not due to impaired microtubule nucleation or microtubule-based transport, as these processes in PCM1-depleted cells were comparable to wild-type (WT) cells, as tested by microtubule regrowth and Golgi-distribution assays (Fig. S3D-F).

**Figure 3.**
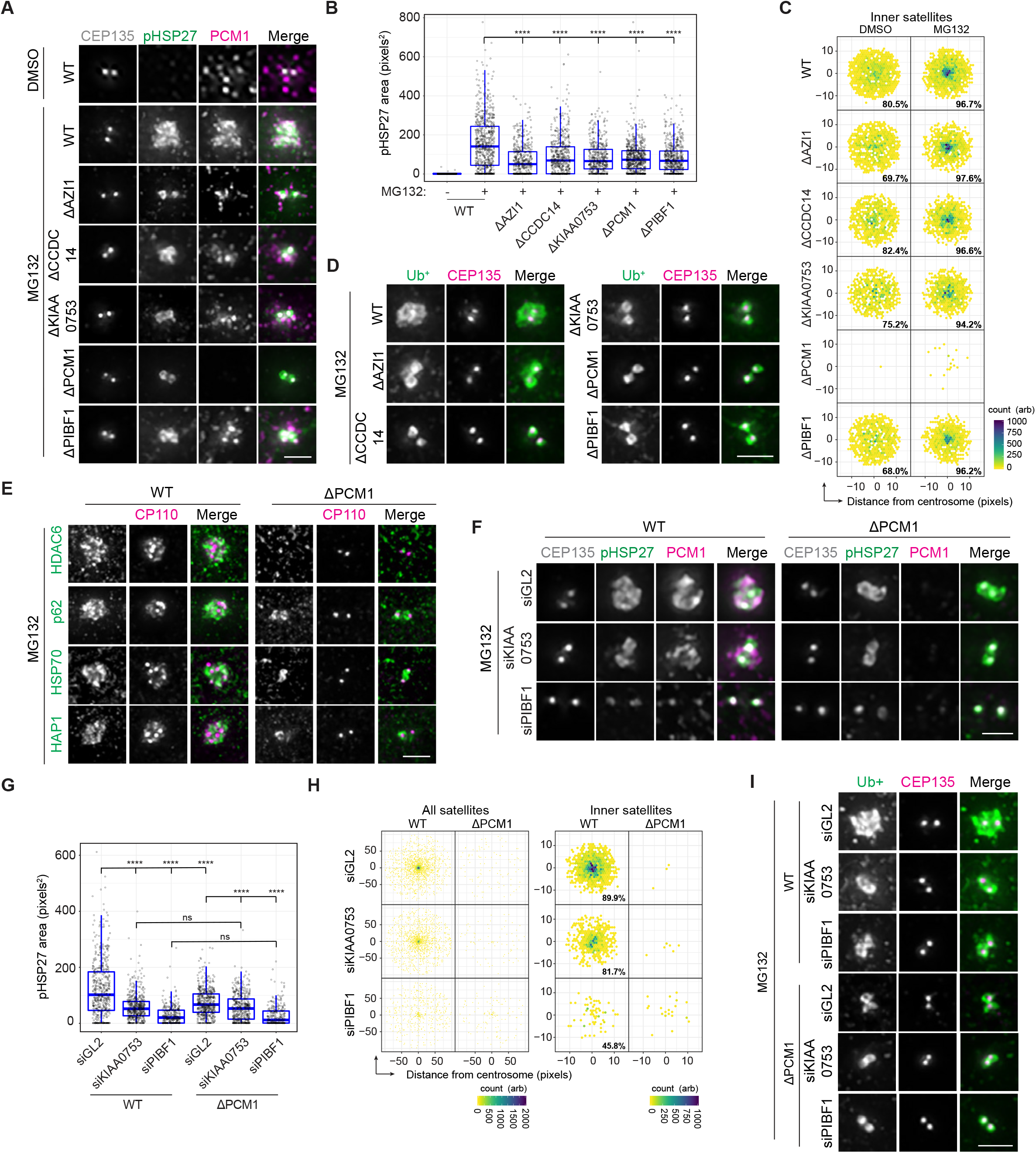
Centriolar satellites are required for aggresome formation. **A**. WT and AZI1, CCDC14, KIAA0753, PCM1 and PIBF1 KO RPE-1 cell lines were treated with MG132 and stained with antibodies against CEP135, pHSP27 and PCM1. **B**. Box-and-whisker plot showing the area occupied by pHSP27 in cells treated as in A. **C**. Intensity maps of PCM1 distribution relative to the centrosome in cells treated as treated in A. The percentage PCM1 signal residing in the defined ‘inner’ region is indicated. **D**. Ub^+^ proteins and CEP135 staining in WT and satellite protein KO cells treated with MG132. **E**. WT and PCM1 KO cells treated with MG132 were stained for CP110 and the indicated protein. **F**. WT and PCM1 KO cells were treated with siRNA against GL2 (control), KIAA0753 or PIBF1 as indicated for 48 hours, then treated with MG132. Cells were stained with antibodies against CEP135, pHSP27 and PCM1. **G**. Box-and- whisker plot showing the area occupied by pHSP27 in cells treated as in F. **H**. Intensity maps of PCM1 distribution relative to the centrosome in cells treated as in F. The percentage PCM1 signal residing in the defined ‘inner’ region is indicated. **I**. Ub^+^ proteins and CEP135 staining in WT and PCM1 KO cells treated as in F. Scale bars: A, D, E, F and I 2 μm. ****p < 0.0001; ns, not significant. See also Figure S3.

Despite lacking satellites, PCM1 KO cells retain a centrosomal pool of AZI1, CCDC14, KIAA0753 and PIBF1 (Fig. S3B). That each KO line is as impaired in aggresome formation as the PCM1-depleted line, suggests that the satellite pool of these proteins is involved in aggresome assembly, rather than their centrosomal fractions. To test this, we sought to knockdown KIAA0753 and PIBF1 by siRNA in WT and PCM1 KO cells. Successful depletion of each protein was confirmed by IF microscopy and immunoblot (Fig. S3G-H), although we observed that KIAA0753 levels were also reduced in WT cells depleted of PIBF1. The distribution of satellites in control and KIAA0753-depleted WT cells was comparable, although depletion of PIBF1 reduced the abundance of satellites around the centrioles (Fig. S3I-J). KIAA0753 and PIBF1 knockdown reduced aggresome formation in WT cells (Fig. 3F-G), and while satellites accumulated in cells depleted of KIAA0753 similar to controls, this was not observed in PIBF1-depleted cells (Fig. 3H). Depletion of PIBF1 had the greatest effect on aggresome assembly, potentially due to the reduction in KIAA0753 abundance and/or the limited movement of satellites into the pericentrosomal region in these cells.

In PCM1 KO cells, KIAA0753 and PIBF1 knockdown modestly reduced aggresome size further, with PIBF1 again having the greatest effect. This suggests that contribution of the centrosomal pool of these proteins to aggresome formation is minor. As PIBF1 depletion did not affect KIAA0753 levels in PCM1 KO cells (Fig. S3H), this supports the notion that the effect of PIBF1 knockdown in WT cells is due to reduced abundance of PIBF1 and not KIAA0753. Upon knockdown of KIAA0753 or PIBF1 in both WT and PCM1-deficient cells, the accumulation of Ub^+^ proteins was reduced (Fig. 3I). Together, these results demonstrate the requirement for intact centriolar satellites in aggresome formation following proteasome inhibition, and suggest an overlapping function of satellite proteins in this process.

### Knockdown of CCNF can induce the formation of aggresome-like structures in the absence of proteasome inhibition

Proteolysis plays a crucial role in the regulation of centriole duplication (Nigg and Holland, 2018) and treatment of U-2 OS cells with the proteasome inhibitor Z-L3VS for 48 hours results in centriole amplification as determined by CETN-GFP imaging (Duensing et al., 2007). Staining for centriole markers typically reveals 2 or 4 foci, corresponding to 2 or 4 centrioles, depending on cell cycle phase. After treating RPE-1 cells with MG132, we observed greater than 4 foci of CETN2, CP110 and CEP97 within the aggresome, although CEP135, SAS6 and GT335 labelling revealed a normal complement of centrioles (Fig. 4A-B). This suggests that the extra foci of CETN2, CP110 and CEP97 in the aggresome do not correspond to supernumerary centrioles. To confirm this, we performed transmission electron microscopy (TEM) on DMSO and MG132 treated cells and did not observe excess centrioles in any of the sections examined (n = 50, Fig. 4C).

**Figure 4.**
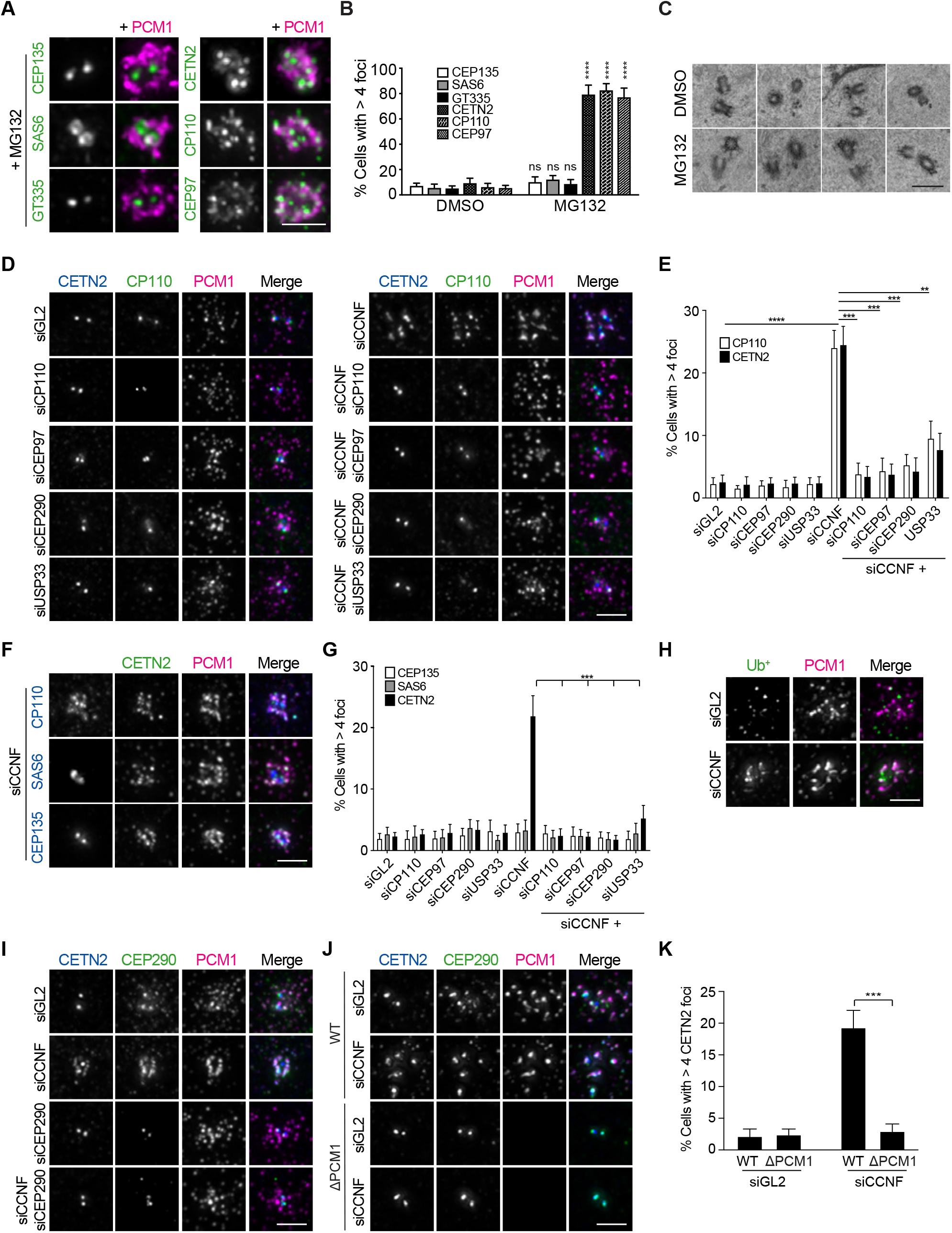
Depletion of CCNF induces the formation of aggresome-like structures without MG132 treatment. **A**. RPE-1 cells treated with MG132 were stained for PCM1 and the indicated protein. **B**. Histogram showing the percentage of cells with greater than 4 foci of CEP135, SAS6, GT335, CETN2, CP110 and CEP97 following MG132 treatment. **C**. TEM images of centrioles from RPE-1 cells treated with DMSO or MG132. **D**. WT cells were treated with siRNAs against GL2 (control) and CCNF, plus CP110, CEP97, CEP290 and USP33, as indicated. After 72 hours, cells were stained for CETN2, CP110 and PCM1. **E**. Histogram of the percentage of cells treated as in D that have more than 4 foci of CETN2 or CP110 per cell. **F**. RPE-1 cells depleted of CCNF for 72 hours were stained for CETN2 and PCM1 along with CP110, SAS6 or CEP135 as indicated. **G**. Histogram of the percentage of cells treated with the indicated siRNAs that have more than 4 foci of CEP135, SAS6 or CETN2 per cell. **H**. RPE-1 cells depleted of GL2 or CCNF were stained for Ub^+^ proteins and PCM1. **I**. RPE-1 cells depleted of GL2, CCNF, CEP290 or CCNF plus CEP290, were stained for CETN2, CEP290 and PCM1. **J**. WT and PCM1 KO cells were depleted of GL2 or CCNF and stained for CETN2, CEP290 and PCM1. **K**. Histogram of the percentage of cells treated as in J that have more than 4 foci of CETN2 per cell. Scale bars: A, D, F, H, I and J 2 μm; C 500 nm. ****p < 0.0001; ***p < 0.001; **p < 0.01; ns, not significant. See also Figure S4.

CP110 is targeted for destruction by Cyclin F (CCNF), with both CCNF knockdown and MG132 treatment stabilizing CP110 levels (D’Angiolella et al., 2010; Li et al., 2013). CCNF depletion has been reported to result in the accumulation of supernumerary centrosomes as scored by the accumulation of excess foci of CETN2, CP110 and γ-tubulin (D’Angiolella et al., 2010). As these three proteins accumulate into aggresomes following proteasome inhibition, and no other centriole markers were utilized in that earlier study, we wondered whether CCNF depletion leads to the formation of aggresome-like structures, rather than extra centrioles. Knockdown of CCNF was confirmed by immunoblot (Fig. S4A) and, in line with previous reports, we recorded ~20% of cells with more than 4 CETN2 or CP110 foci following CCNF depletion (Fig. 4D-E). In addition to co-localizing with CP110, these supernumerary CETN2 foci also contained PCM1 (Fig. 4D).

Knockdown of CP110 successfully reduced its total abundance in cells, while a more stable fraction persisted at the centrioles (Figs. 4D and S4A). As previously reported, the accumulation of excess CETN2 foci in cells depleted of CCNF required CP110 and its DUB USP33 (D’Angiolella et al., 2010; Li et al., 2013), and we found it also depended upon CEP97 and CEP290 (Fig. 4D-E). Furthermore, depletion of CP110 and CEP290 also blocked MG132-induced accumulation of CETN2, CP110 and PCM1 (Fig. S4B-C). To assess the number of centrioles in cells depleted of CCNF, we looked to SAS6 and CEP135, two centriole markers that do not accumulate into aggresomes during proteasome inhibition. Despite CCNF-depletion leading to cells containing extra foci of CETN2, there was no accumulation of supernumerary SAS6 or CEP135 foci (Fig. 4F-G). This indicates that knockdown of CCNF does not lead to the formation of extra centrioles, thereby challenging previous reports.

Further support for induction of aggresome-like structures by CCNF-depletion was provided by staining for Ub^+^ proteins. In control cells, small Ub^+^ foci were seen in the vicinity of centriolar satellites, but they did not co-localize with PCM1. However, in CCNF-depleted cells, Ub^+^ proteins co-localize with clustered satellites (Fig. 4H). Similarly, CEP290 also co-localized with the extra CETN2 foci in cells depleted of CCNF (Fig. 4I). Depletion of CEP290 saw its removal from satellites while still being retained on the centrioles. Co-depletion of CEP290 with CCNF looked comparable to CEP290 depletion alone, with CEP290 restricted to the centrioles and no extra CETN2 foci being formed (Fig. 4I). Accordingly, CEP290 remained restricted to CETN2 positive-centrioles in cells in which MG132-induced aggresome formation had been blocked by CEP290 depletion (Fig. S4C). Furthermore, PCM1 KO cells were unable to form additional CETN2/CEP290/PCM1 positive foci upon depletion of CCNF, indicating that satellites play a role in this process (Fig. 4J-K), while overexpression of CCNF prevented aggresome formation in MG132-treated cells (Fig. S4D-E). Together, these data support that dysregulation of CCNF leads to the formation of aggresome-like structures through CP110, CEP97, CEP290 and PCM1, most likely via its known role as an SCF substrate recognition unit.

### The CP110-CEP97-CEP290 module is required for aggresome formation

Having observed a block to the accumulation of CETN2, CP110, CEP290 and PCM1 in MG132-treated cells when either CP110 or CEP290 were depleted (Fig. S4B-C), we sought to measure aggresome formation in these cells using our quantification pipeline. We also included cells depleted of CEP97, CCNF and USP33 in this analysis. Knockdown of CP110, CEP97, CEP290 and USP33 reduced the capacity for aggresome formation below that of control and CCNF-depleted cells (Fig. 5A-B). Strikingly, knockdown of CP110 had the greatest effect, and caused mis-localization of pHSP27 to actin filaments (Fig. S5A). An RNAi-resistant version of CP110 was recruited into the aggresome when expressed in cells, and was able to rescue aggresome formation in CP110-depleted cells (Fig. S5B-E). Conversely, when CEP290 transfected cells were treated with MG132, multiple aggresome-like structures containing pHSP27 and PCM1 were observed throughout the cytoplasm (Fig. S5F), suggesting CEP290 can drive the assembly of aggresomes at ectopic sites.

**Figure 5.**
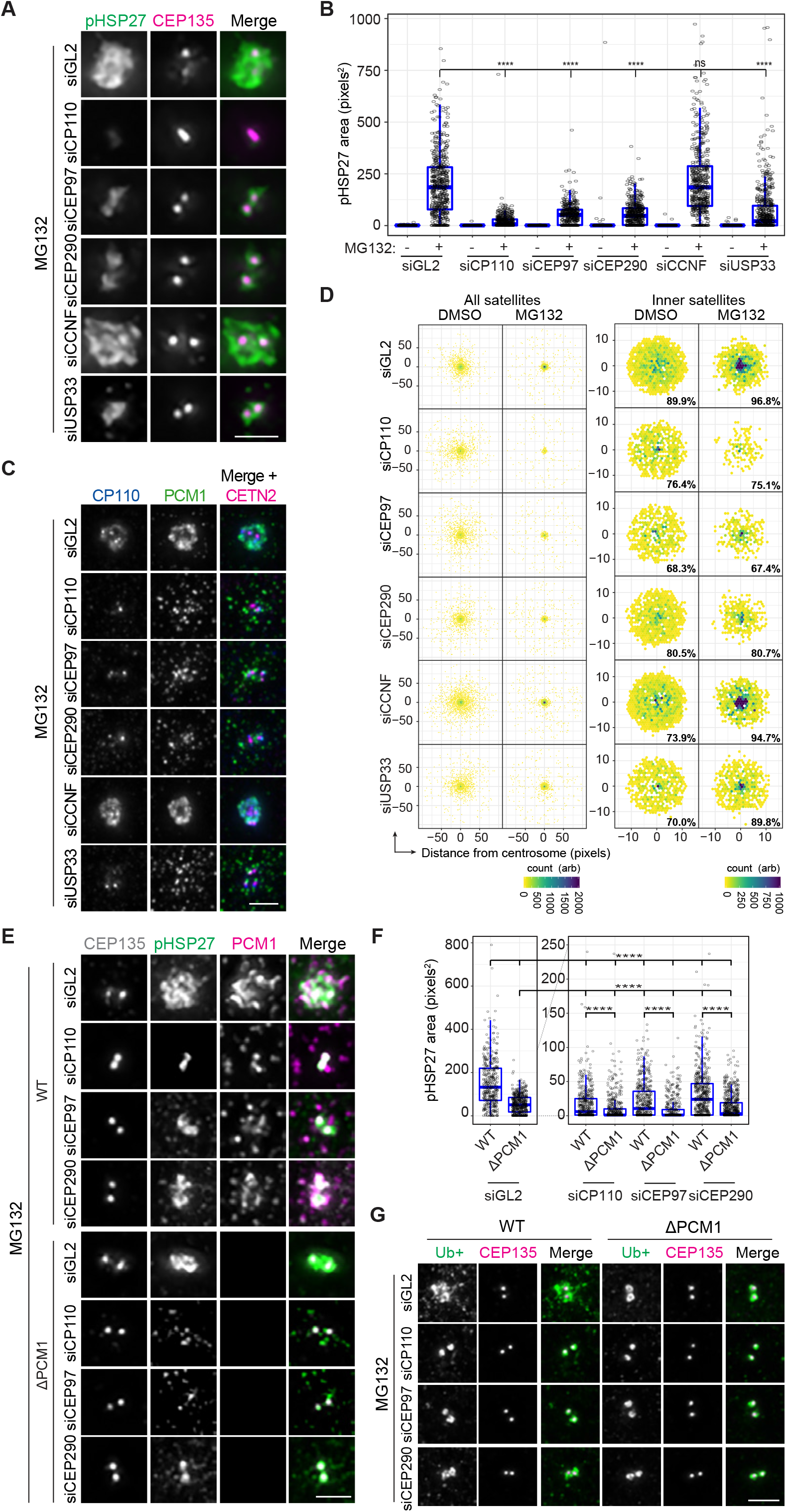
A CP110-CEP97-CEP290 module is required for aggresome formation. **A**. RPE-1 cells depleted of GL2, CP110, CEP97, CEP290, CCNF, or USP33 by siRNA for 48 hours then treated with MG132 were stained for pHSP27 and CEP135. **B**. Box-and-whisker plot showing the area occupied by pHSP27 in cells treated as in A. **C**. Cells treated as in A were stained for CP110 and PCM1. **D**. Intensity maps of PCM1 distribution relative to the centrosome in cells treated as in A. The percentage PCM1 signal residing in the defined ‘inner’ region is indicated. **E**. WT and PCM1 KO cells were depleted of GL2, CP110, CEP97 or CEP290 by siRNA for 48 hours then treated with MG132 and stained for CEP135, pHSP27 and PCM1. **F**. Box-and-whisker plot showing the area occupied by pHSP27 in cells treated as in E. Data from the same experiment are displayed on side-by-side plots with different y-axis ranges to emphasize the difference between the WT and PCM1 KO cells. **G**. Cells treated as in E were stained for Ub^+^ proteins and CEP135. Scale bars: A, C, E and G 2 μm. ****p < 0.0001; ns, not significant. See also Figure S5.

Examination of satellite distribution in cells depleted of CP110-CEP97-CEP290 module components revealed an accumulation around the centrioles in control, CCNF and USP33 depleted cells, that was reduced in cells depleted of CP110, CEP97 and CEP290 (Fig. 5C-D). To determine whether the effects of CP110, CEP97 and CEP290 knockdown on aggresome formation operate through satellites, we quantified pHSP27 in PCM1 KO cells depleted of these proteins. For each protein, their knockdown in PCM1-deficient cells reduced aggresome formation further, with the depletion of CP110, CEP97 or CEP290 in the absence of PCM1 having an additive effect on aggresome assembly (Fig. 5E-F). This suggests that satellites are required for a fraction of the recruitment of pHSP27 to the centrioles in the absence of the CP110, CEP97 or CEP290, and that CP110, CEP97 and CEP290 are responsible for the accumulation of a proportion of pHSP27 in the absence of satellites. The recruitment of Ub^+^ proteins to the centrosomal region was reduced accordingly in cells depleted of CP110, CEP97 or CEP290 (Fig. 5G). Together, these data assign a novel role to the CP110-CEP97-CEP290 module in aggresome formation.

### Centriolar satellites and the CP110-CEP97-CEP290 module are required for aggresome formation in senescent cells and a Huntington’s disease model

Ageing is the primary risk factor for a range of diseases, including neurodegenerative disorders (Hou et al., 2019). Decreased proteasomal activity in senescent cells results in the accumulation of protein aggregates, which correlate with age-related diseases (Cuanalo-Contreras et al., 2013; Fernández-Cruz and Reynaud, 2020; López-Otín et al., 2013; Saez and Vilchez, 2014). As aggresome formation serves a protective function through the accumulation of protein aggregates into a single location, we wondered whether senescent cells are able to utilize the aggresome pathway. Somatic cells in culture undergo replicative senescence after a finite number of divisions (Hayflick, 1965). To obtain senescent cells, we grew low passage number primary Human Foreskin Fibroblasts (HFF-1) for an extended period of time of at least 140 days. Induction of cellular senescence was confirmed by the presence of senescence-associated ß-galactosidase (SA-ß-gal) activity (Fig. S6A; Dimri et al., 1995). Increased expression of p53 and p21 was also observed, indicative of a senescent population (Fig. S6B; Rufini et al., 2019). When treated with MG132, the majority of the senescent population were unable to form aggresomes, which correlated with cells being negative for the proliferation marker Ki67 (Fig. 6A-B). Instead of accumulating into an aggresome, Ub^+^ proteins were observed as smaller aggregates throughout the cytoplasm in senescent cells (Fig. S6C).

**Figure 6.**
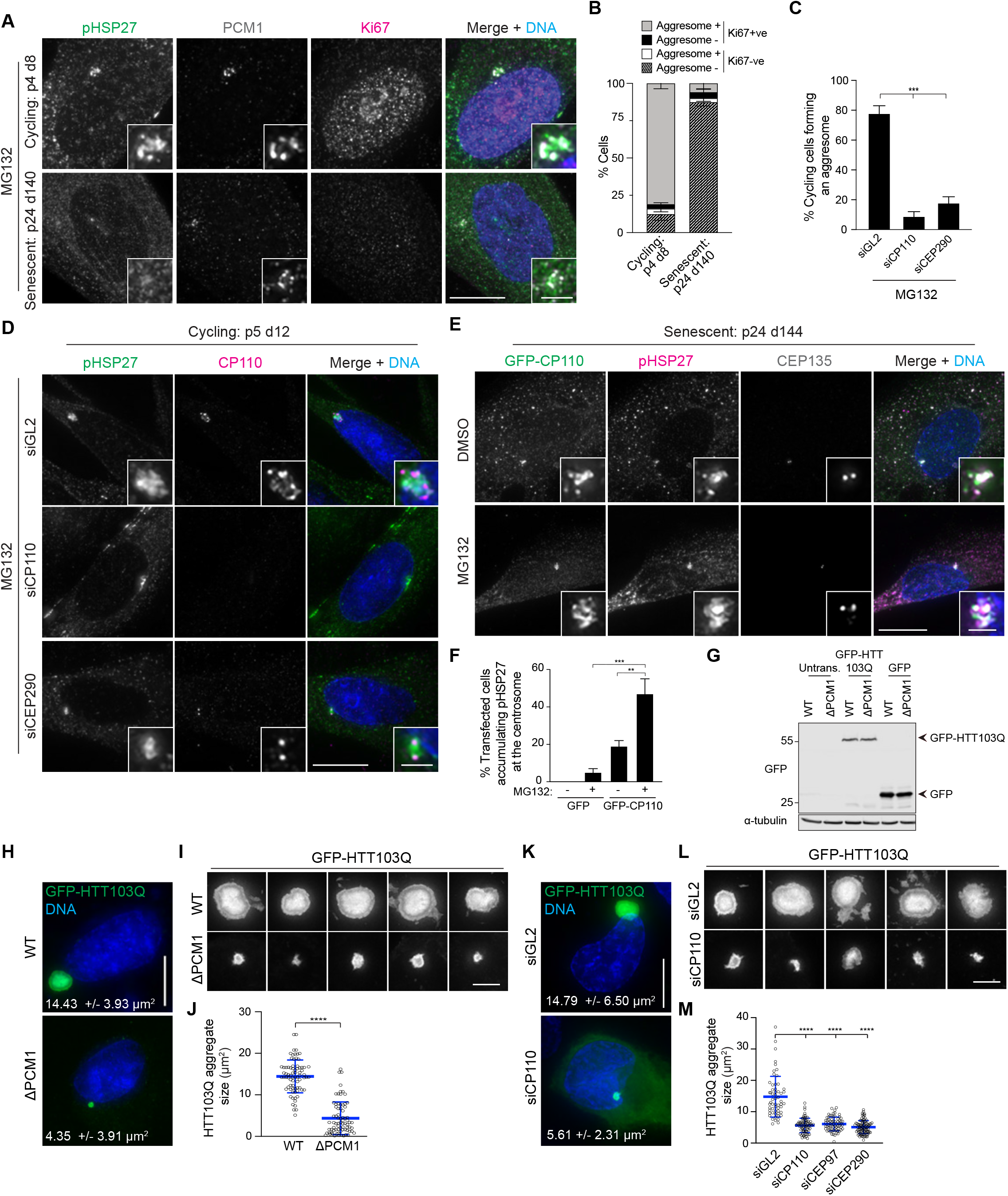
Senescent cells have a reduced capacity to form aggresomes and HTT-polyQ aggregation requires centriolar satellites and the CP110-CEP97-CEP290 module. **A**. Cycling and senescent HFF-1 cells were treated with MG132 and stained for pHSP27, PCM1, Ki67 and DNA (DAPI). **B**. Quantitation of aggresome formation in Ki67 positive and negative cycling and senescent HFF-1 cells. **C**. Quantitation of aggresome formation in cycling HFF-1 cells depleted of GL2 (control), CP110 or CEP290 and treated with MG132. **D**. Cells treated as in C were stained for pHSP27, CP110 and DNA (DAPI). **E**. Senescent HFF-1 cells were transiently transfected with GFP-CP110 for 48 hours before treatment with MG132. Cells were stained for pHSP27, CEP135 and DNA (DAPI). **F**. Histogram of the percentage of senescent HFF-1 cells transfected with GFP or GFP-CP110 that were able to form an aggresome after MG132 treatment. **G**. Immunoblot of extracts from WT and PCM1 KO RPE-1 cells transiently transfected with GFP or GFP-HTT103Q and probed for GFP. α-tubulin was used as a loading control. **H**. WT and PCM1 KO RPE-1 cells were transiently transfected with GFP-HTT103Q for 24 hours, then fixed and the DNA stained with DAPI. The mean aggregate size is indicated. **I**. Examples of GFP-HTT103Q aggregates forming in WT and PCM1 KO RPE-1cells. **J**. Box-and-whisker plot of GFP-HTT103Q aggregate size in WT and PCM1 KO cells. **K**. WT cells were treated with siRNAs against GL2 and CP110, then transiently transfected with GFP-HTT103Q for 24 hours, then fixed and the DNA stained with DAPI. The mean aggregate size is indicated. **L**. Examples of GFP-HTT103Q aggregates forming in WT cells depleted of GL2 (control) or CP110. **M**. Box-and-whisker plot of GFP-HTT103Q aggregate size in WT cells depleted of GL2 (control), CP110, CEP97 and CEP290. Scale bars: A, D, E, H and K 10 μm, insets A, D and E 2 μm; I and L 2 μm. ****p < 0.0001; ***p < 0.001; **p < 0.01. See also Figure S6.

We noted decreased CP110 levels in senescent cells, as compared to their cycling counterparts (Fig. S6B), which was in line with a previous study (Breslin et al., 2014). As we have established a role for CP110 in aggresome formation, we questioned whether the amounts of CP110 in senescent cells were limiting aggresome assembly in these cells. First, depletion of CP110 and CEP290 in cycling HFF-1 cells recapitulated our earlier findings, as knockdown of either protein reduced the percentage of cells that could form an aggresome (Fig. 6C-D). Conversely, expression of CP110 in senescent cells was able to rescue aggresome formation (Fig. 6E-F). Strikingly, CP110 expression was able to induce the accumulation of pHSP27 at the centrosome in a proportion of cells even in the absence of proteasome inhibition. Together, these results demonstrate that senescent cells have a reduced capacity to form aggresomes compared to cycling cells and that CP110 levels are a limiting factor to aggresome formation.

The enrichment of HTT, with ubiquitin, in aggresome-like structures is a pathological feature of HD (Olzmann et al., 2008). As HTT has been shown to interact with PCM1 through HAP1 (Keryer et al., 2011), we asked whether satellites, or the CP110-CEP97-CEP290 module, are required for the accumulation of a HTT fusion protein containing 103 polyQ repeats (GFP-HTT103Q) into inclusions. Upon transfecting GFP-HTT103Q for 24 hours, we found that cells predominantly contained a single large GFP-positive inclusion to which pHSP27 and Ub^+^ localized (Fig. S6D-E), suggesting that GFP-HTT103Q inclusions might possess similar properties to aggresomes. To test a requirement for satellites in HTT-polyQ inclusion assembly, we transfected GFP-HTT103Q into WT and PCM1 KO RPE-1 cells. Equivalent levels of expression were achieved in each cell line (Fig. 6G). Measurement of GFP-HTT103Q inclusions in each line revealed that they were restricted in size to 4.35 +/− 3.91 μm^2^ in PCM1 KO cells, compared to 14.43 +/− 3.93 μm^2^ in WT cells (Fig. 6H-J). Inclusion size was similarly restricted in cells depleted of CP110, CEP97 or CEP290 (Fig. 6K-M). Therefore, these results support a role for satellites and the CP110-CEP97-CEP290 module in the pathologically relevant aggregation of proteins.

## DISCUSSION

Here, we have described the requirement for the CP110-CEP97-CEP290 module and centriolar satellites in aggresome formation, with high-resolution quantitative analysis of aggresome assembly allowing us to delineate discrete steps in the aggresome pathway (Fig. 7). In the absence of active protein translation, aggresome formation was completely blocked. The initial recruitment of pHSP27 to the centrioles ‘seeds’ aggresome assembly. This foundation of pHSP27 expands in a manner that depends first on the CP110-CEP97-CEP290 module and then centriolar satellites. Finally, further recruitment to the aggresome requires satellites, microtubules and active HDAC6.

**Figure 7.**
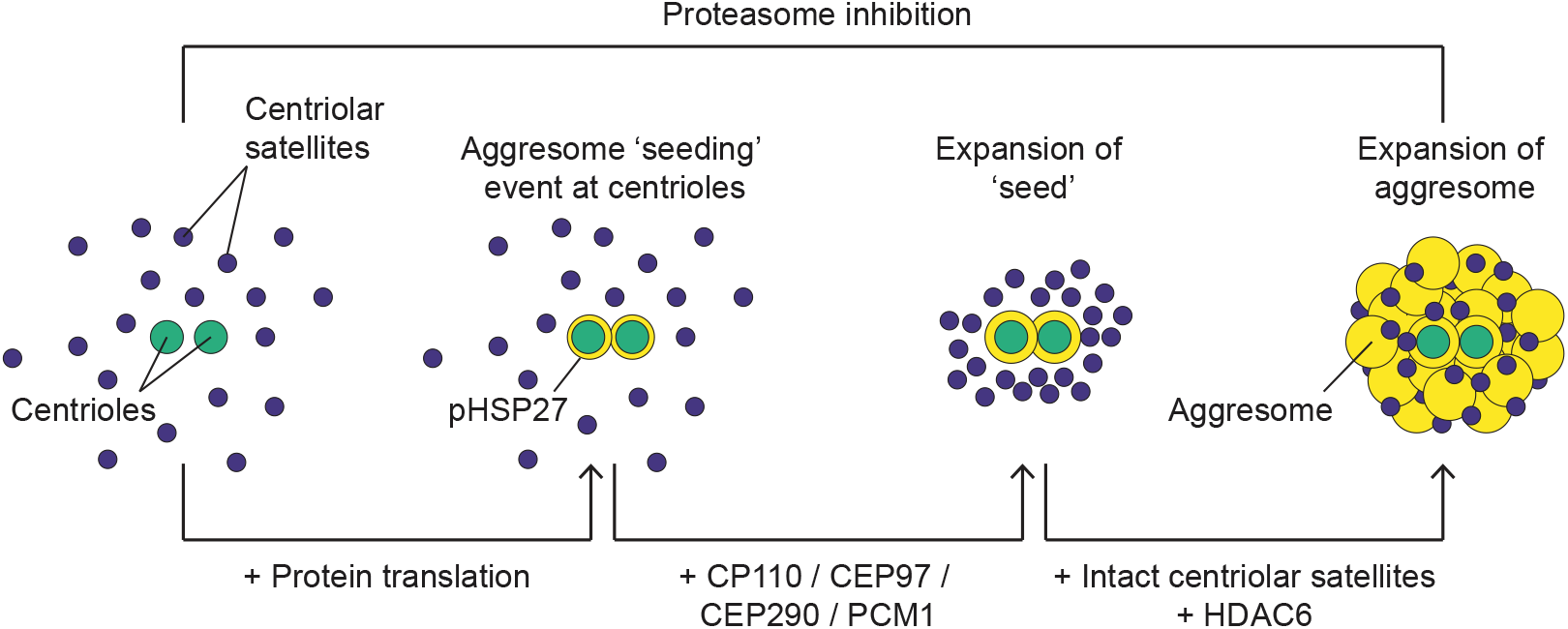
Schematic representation of the aggresome assembly pathway. The earliest event in aggresome formation at the centrosome is the appearance of pHSP27 on the centrioles, which required active protein translation. This acts as a seeding event, laying a foundation for recruitment of proteins that will become incorporated into the aggresome. The first step in the expansion of pHSP27 requires CP110, CEP97, CEP290 and PCM1. Further expansion depends on intact centriolar satellites and HDAC6.

Knockdown or overexpression of individual satellite proteins has previously been shown to promote satellite aggregation (Hori et al., 2015; Kim et al., 2008; 2004; Stowe et al., 2012), suggesting satellites play a finely tuned role in maintaining proteostasis. In support of this, wide-ranging changes in the cellular proteome in cells devoid of satellites confirmed them as regulators of global proteostasis (Odabasi et al., 2019). Our findings place satellites at the nexus between centrosomes and the proteostasis networks in the response to proteotoxic stress. Sequestration of proteins for destruction at the centre of the microtubule network allows proteins to be transported from throughout the cytoplasm to a central location for further processing. Satellites functioning as conduits for the movement of aggregated proteins to the pericentriolar region is the simplest explanation for their role in aggresome assembly. Satellites associating with ubiquitinated aggregates in the cytoplasm during proteasome inhibition supports their early recruitment to these structures. Intriguingly, satellites increased in density around the centrosome even when aggresome formation was blocked by inhibition of HDAC6, suggesting that their movement to this region does not depend on interactions with aggresomal cargo. Interestingly, AZI1 possesses a putative HDAC-interacting domain, although an interaction with HDAC6 has yet to be determined (Wilkinson et al., 2009). Identification of the molecular linker that couples aggregated proteins to satellites for their transit to the aggresome will be the focus of future studies.

The UPS and autophagy are the main protein degradation pathways in the cell, and they are demonstrably interconnected as disruption of one causes upregulation of the other (Kocaturk and Gozuacik, 2018; Liu et al., 2016; Selimovic et al., 2013; Wang et al., 2014; Wu et al., 2008). Satellites regulate the stability of proteins via both the UPS and autophagy (Joachim et al., 2017; Wang et al., 2016). Indeed, depletion of satellite proteins OFD1 or BBS4 led to deficiencies in the proteasome-degradation pathway (Gerdes et al., 2007; Liu et al., 2014). UPS activity and autophagy also regulate satellite composition: the E3 ligase MIB1 ubiquitylates PCM1 and AZI1 to promote their degradation (Wang et al., 2016), while their stability is promoted by the DUBs CYLD and USP9X (Douanne et al., 2019; Han et al., 2019; Li et al., 2017; Wang et al., 2017); and degradation of PCM1 through selective autophagy regulates satellite turnover (Holdgaard et al., 2019). Together, these findings place satellites at the intersection of the two degradation pathways. As aggresomes are cleared by selective autophagy (Choi et al., 2020; Fortun et al., 2003; Hao et al., 2013), the requirement of satellites for aggresome formation in cells with inhibited proteasomes thereby strengthens this position.

Observing that depletion of CCNF led to the formation of aggresome-like structures, rather than centriole overduplication as previously reported (D’Angiolella et al., 2010), allowed us to establish a role for the CP110-CEP97-CEP290 module at one of the earliest points of aggresome assembly at the centrosome. Consequently, this assigns a new function to CP110 and its partners beyond the control of centriole length. The requirement for the CP110 module for the formation of a ring of pHSP27 around the centrioles, suggests that this is an initial seeding event that acts as a foundation for the assembly of aggresomal particles into a single structure. Not only does this imply that transport of aggregates to the centrosome is highly regulated, but also that the building of an aggresome from these components is controlled, to allow an ordered structure to form. The images we present here support the notion that the aggresome has higher-order structure. Indeed, early TEM revealed that multiple particles loosely associate with each other within an aggresome, rather than coalescing into a single large aggregate (Garcia-Mata et al., 2002; García-Mata et al., 1999; Johnston et al., 1998). Aggregated proteins tend to be sticky due to misfolding exposing hydrophobic residues. Ordered structure within the aggresome would mitigate the presence of these residues, preventing further aggregation and interaction with other cellular components (Drummond, 2012; Garcia-Mata et al., 2002), providing additional support for the protective role the aggresome plays.

Ageing is the biggest risk factor associated with most neurodegenerative disorders (Hou et al., 2019), with widespread protein aggregation a common feature of aged cells. Collapse of proteostasis is a driver of age-related aggregate formation (Santra et al., 2019), occurring through the loss of protein-quality-control and decreased proteasomal activity (Keller et al., 2000). We found that senescent cells have a reduced capacity to form aggresomes, which was due to limiting amounts of CP110. Further to this, we found that satellites and the CP110-CEP97-CEP290 module were required for the aggregation of polyglutamine-containing HTT. Despite protein aggregates being a hallmark of neurodegenerative disorders, the contribution of these inclusions to the pathologies of these diseases remains unclear. Primary cilium structure and function is altered in HD (Kaliszewski et al., 2015), with cilia being crucial to the normal functioning of neurons (Lee and Gleeson, 2010). Interestingly, HAP1, which binds to HTT in association with the polyglutamine repeat (Li et al., 1998), interacts with PCM1 (Engelender et al., 1997), and expression of pathogenic polyglutamine expansions leads to the accumulation of PCM1 at the centrosome and altered cilium structure (Keryer et al., 2011). Furthermore, aggresomes of α-synuclein, increased levels of which are sufficient to cause Parkinson’s disease, have been shown to inhibit ciliogenesis and centrosome function (Iqbal et al., 2020). These findings suggest that there is an intricate link between protein aggregates and cilium function in neurodegeneration. Given the emergent connection between satellites and aggregate processing, and the requirement for satellites in cilia formation and function (Odabasi et al., 2019; Wang et al., 2016), future studies to elucidate the role of satellites in these processes will provide valuable insight into the pathophysiology of neurodegenerative diseases and may help to illuminate potential therapeutic avenues.

## ACKNOWLEDGMENTS

We thank members of the Pelletier Lab for their scientific feedback during the project. We are grateful to G.G. Hesketh for sharing the GFP-HTT-polyQ construct. We thank R. Buijs for help generating U-2 OS STIL KO cells. We are grateful to S. Cheung and K. Lane for proofreading the manuscript. S.L.P. was funded by a European Union Horizon 2020 Marie Skłodowska-Curie Global Fellowship (No. 702601). This work was funded by a CIHR Foundation Grant, Canada Research Chair, and the Krembil Foundation to L.P.

## AUTHOR CONTRIBUTIONS

Conceptualization, S.L.P., C.G.M., and L.P.; Methodology, S.L.P. and J.T.; Formal Analysis, S.L.P. and J.T.; Investigation, S.L.P.; Resources, J.T. and L.G.; Writing – Original Draft, S.L.P. and L.P.; Writing – Review & Editing, S.L.P, J.T., C.G.M., and L.P.; Visualization, S.L.P. and J.T.; Supervision, C.G.M and L.P.; Funding Acquisition, S.L.P and L.P.

## MATERIALS AND METHODS

### Cell culture and drug treatments

All cell lines were cultured in a 5% CO_2_ humidified atmosphere at 37°C. hTERT RPE-1 (female, human epithelial cells immortalized with hTERT), A-375 (female, human malignant melanoma epithelial), BJ-5ta (male, human fibroblasts immortalized with hTERT), HFF-1 (male, human primary fibroblasts), HeLa (female, human adenocarcinoma epithelial) and U-2 OS (female, human osteosarcoma epithelial) cells from ATCC were grown in Dulbecco’s Modified Eagle’s Medium (DMEM; Life Technologies) supplemented with 10% v/v fetal bovine serum. To inhibit the proteasome, cells were treated with 10 μM MG132 (Millipore-Sigma) or 1 μM bortezomib (Santa Cruz Biotechnology) for 5 hours. To disrupt microtubules, cells were treated with nocodazole (Millipore-Sigma) at 10 μM or taxol (paclitaxel; Millipore-Sigma) at 5 μM for 2 hours before the addition of MG132. Cycloheximide (Enzo Life Sciences) was used to inhibit protein translation at 5 μg/ml. ACY-1215 and ACY-738 (Selleck Chemicals) at 50 μM each were used to inhibit HDAC6. As control, cells were treated with vehicle (DMSO) alone. For microtubule regrowth assays, cells were treated with nocodazole for 5 hours before drug washout with pre-warmed media for 60 seconds. To verify senescent populations, senescence associated β-galactosidase activity was assayed using a Senescence Detection Kit (Abcam) according to the manufacturer’s instructions.

### Generation of knockout cell lines

To generate hTERT RPE-1 AZI1/CEP131, CCDC14, KIAA0753 and PIBF1/CEP90 knockout cell lines, guide RNA targeting the respective gene (see Table S1) was selected and transcribed *in vitro* before being transfected into wild-type hTERT RPE-1 cells constitutively expressing Cas9 using Lipofectamine RNAiMAX (Invitrogen) according to the manufacturer’s instructions. After 5 days, the transfected cells were diluted so that single clones could subsequently be isolated. Gene disruption was confirmed by PCR amplification of genomic DNA with the primers detailed in Table S1 and sequence analysis using ICE (Inference of CRISPR Edits, Synthego; Table S1). Loss of signal of the respective protein was demonstrated by immunoblot and immunofluorescence microscopy.

### RNA-mediated interference

All siRNA transfections were performed using Lipofectamine RNAiMAX (Invitrogen) according to the manufacturer’s instructions. Details of siRNA oligos utilized in this study are provided in Table S2. hTERT RPE-1 cells were transfected with 20 nM (final concentration) of the respective siRNA for 48 or 72 hours, as indicated. Effective knockdown was confirmed by immunoblot and/or immunofluorescence microscopy. An RNAi-resistant mutant of CP110 was generated using site-directed mutagenesis and the primers detailed in Table S2. To rescue the CP110 RNAi phenotype, cells were transfected with 1 μg of DNA using Lipofectamine 3000 (Invitrogen) 24 hours post-siRNA transfection. After a further 24 hours, cells were treated with MG132 for 5 hours.

### Transient transfections

Transfections were performed using Lipofectamine 3000 (Invitrogen) according to the manufacturer’s instructions. Briefly, 1 μg plasmid DNA was complexed with Lipofectamine 3000 in serum-free Opti-MEM and added to cells at 70-80% confluency. 24 hours after transfection, cells were treated with MG132 for 5 hours before analysis by immunofluorescence microscopy.

### Immunofluorescence microscopy

Details about the primary and secondary antibodies used for immunofluorescence microscopy are provided in Table S2. For immunofluorescence microscopy, cells were fixed with ice-cold methanol for 10 minutes at −20°C, with the exception of the cells presented in Fig. S5A which were fixed with 4% paraformaldehyde for 5 minutes at room temperature before being permeabilized with methanol for 2 minutes. All cells were blocked with 2% bovine serum albumin in PBS for 10 minutes, then incubated with primary antibody in blocking solution for 50 minutes at room temperature. After washing with PBS, cells were incubated with fluorophore conjugated secondary antibodies (Table S2) and DAPI (0.1 μg/ml) in blocking solution for 50 minutes at room temperature. After washing with PBS, coverslips were mounted onto glass slides using ProLong Gold Antifade (Molecular Probes). Cells were imaged using a DeltaVision Elite high-resolution imaging system equipped with a sCMOS 2048 x 2048 pixel camera (GE Healthcare). Z-stacks (0.2 μm) were collected using a 60x 1.42 NA plan apochromat oil-immersion objective (Olympus) and deconvolved using softWoRx (v6.0, GE Healthcare). Images are shown as maximum intensity projections (pixel size 0.1064 μm). The images presented in Figures 1F and S1E were captured using a Nikon A1R-HD25 Point Scanning Confocal equipped with a LU-N4 laser unit, A1-DUG-2 GaAsP multi detector unit and Photometrics Prime95B 25 mm ultra-high sensitivity sCMOS camera. Z-stacks (0.08 μm) were collected using a 60x 1.4 NA plan apochromat lambda oil-immersion objective (Nikon), 16x averaging, 4x zoom, and resonant scanning, and deconvolved in NIS-Elements (Nikon). Selected sections from z-stacks are shown.

### Immunoblotting

Details about the primary and secondary antibodies used for immunoblotting are provided in Table S2. For immunoblotting, total cell lysates were collected in Laemmli buffer and treated with benzonase nuclease (Millipore-Sigma). Proteins were separated on SDS–PAGE gels and transferred to PVDF membrane (Amersham Hybond P, Cytiva). Membranes were incubated with primary antibodies in TBST (TBS, 0.1% Tween-20) in 5% w/v milk powder (BioShop) at 4°C overnight. Following washing in TBST, blots were incubated with IRDye conjugated secondary antibodies (LI-COR) for 1 hour at room temperature. Blots were then washed 3x in TBST and 1x in TBS, before being imaged on a LI-COR Odyssey CLx Infrared Imager.

### Automated quantitative image analysis

Images were analyzed using CellProfiler 3.0 (cellprofiler.org) and R (r-project.org). For pHSP27 analysis, centrosomes were identified using the CEP135 or CETN2 signal. Foci closer than 8 pixels were merged and all centrosomes identified were shrunk to a single pixel. The aggresome area was found using the centrosome coordinates as seed points and expanding outward on the pHSP27 channel using the Watershed algorithm. The lower quartile of each pHSP27 image was used as a background subtraction value before the total pHSP27 intensity in the aggresome area was determined. The results from two independent experiments each using at least 291 cells were pooled and plotted. Boxplots show the median, upper and lower quartiles. The data were analyzed using a Kruskal-Wallis ANOVA to determine if at least one distribution was statistically different from the others. A post-hoc Dunn test was then performed to assess pairwise differences and calculate p-values. For PCM1 analysis, the same images were analyzed and centrosomes identified as above. The centrosome points were expanded 13 pixels to define the ‘inner PCM’ area that approximates the average size of an aggresome. The ‘inner’ region was further expanded 100 pixels to create an annulus defining the ‘outer PCM’ area that generally encompassed the entire cell and were non-overlapping between cells. The ‘inner’ satellites were segmented using an adaptive Otsu method and the ‘outer’ satellites segmented with a global Otsu algorithm. The (x,y) position for every satellite was calculated relative to the cell centrosome position. We assumed that the cells were rotationally similar and overlaid all cells in each condition centred on the centrosome. The PCM1 signal was background subtracted using the lower quartile of each image and further subtracted based on PCM1 knockout images to correct for spurious objects detected in the absence of PCM1. The centroid of each satellite was weighted by the total corrected PCM1 intensity of the object and visualized using a hexbin plot normalized to the total number of cells and replicates. On the ‘inner’ satellite plots, the percentage of total PCM1 intensity within this pericentrosomal region is displayed.

### Statistical methods

Quantitative data in Figures 1C, S1C, 4B, 4E, 4G, 4K, S4E, S5E, 6B, 6C, and 6F represent mean and SD of three independent experiments in which 100 – 200 cells were quantified per condition. HTT103Q aggregate size in Figures 6J and 6M was measured using ImageJ and maximum intensity projections from at least 50 cells per condition captured from two independent experiments. Statistical analyses were performed using two-tailed unpaired Student’s *t*-tests and significance was assumed by p < 0.05. Individual p values are indicated in Figure legends and are defined as: ****p < 0.0001; ***p < 0.001; **p < 0.01; ns, not significant.

### Transmission electron microscopy

For thin-section TEM, hTERT RPE-1 cells were grown in 10 cm dishes and treated with DMSO or MG132 for 5 hours. Cells were then pelleted and washed twice with PBS, before primary fixation in 2% glutaraldehyde and 2% paraformaldehyde in 0.1 M sodium cacodylate buffer overnight at 4°C, and secondary fixation in 2% osmium tetroxide. Samples were dehydrated through an ethanol gradient, followed by propylene oxide, and embedded in EMbed 812 resin (Electron Microscopy Sciences). Ultra-thin sections were cut on an RMC MT6000 ultramicrotome and stained with 2% uranyl acetate in 70% methanol and aqueous lead citrate. Sections were viewed on FEI Tecnai 20 transmission electron microscope.

## SUPPLEMENTAL FIGURE LEGENDS

**Figure S1.**
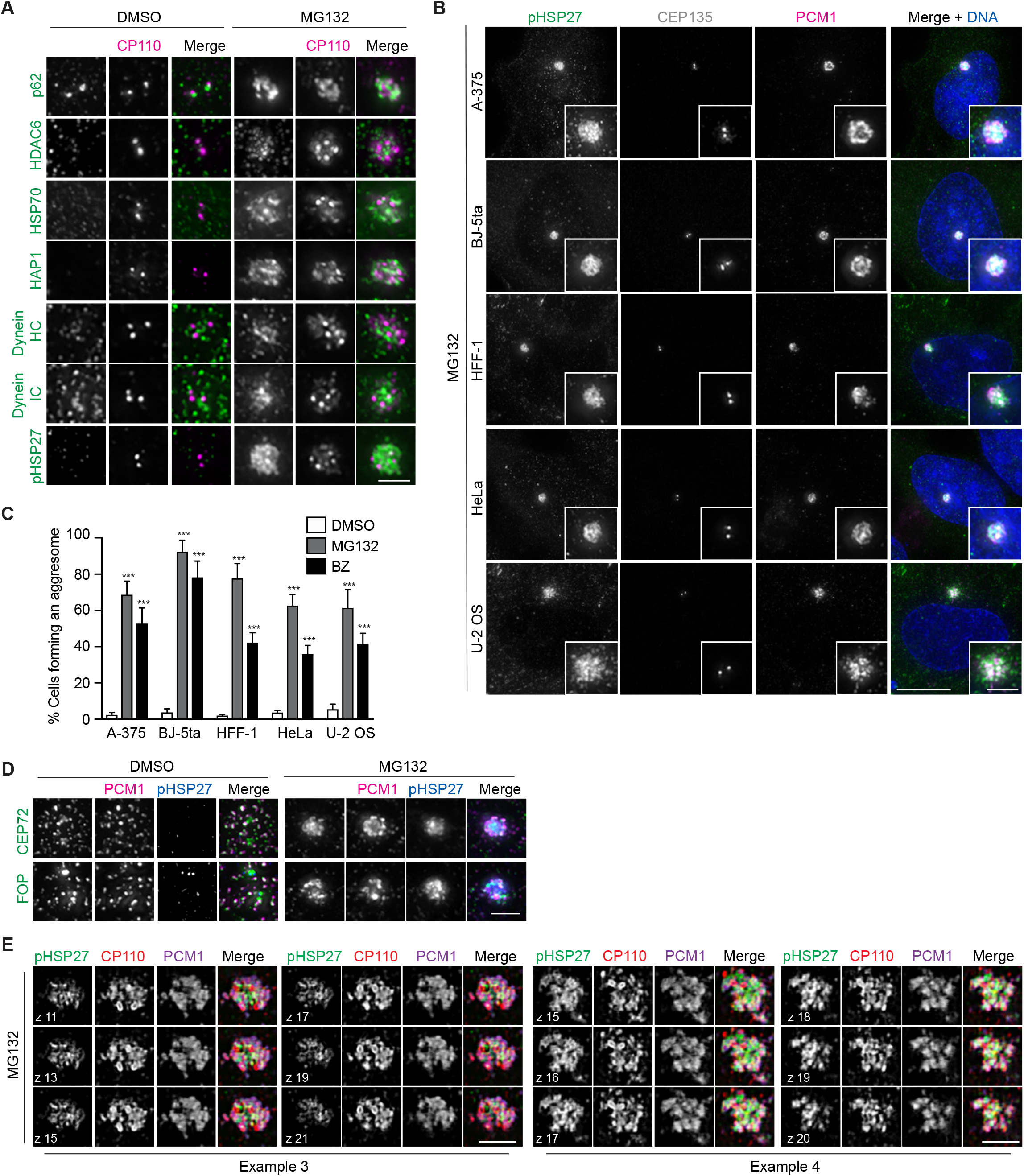
Centrosomal proteins localize to the aggresome upon proteasome inhibition, related to Figure 1. **A**. RPE-1 cells treated with DMSO or MG132 were stained with antibodies against CP110 and the indicated protein. **B.** The indicated cell lines were treated with MG132 and stained with antibodies against pHSP27, CEP135 and PCM1. DNA was stained with DAPI. **C**. Histogram showing the percentage of cells forming an aggresome in DMSO, MG132 and BZ treated cells, as indicated. **D**. RPE-1 cells treated with DMSO or MG132 for 5 hours were stained with antibodies against PCM1, pHSP27 and the indicated protein. **E**. Higher resolution images of aggresomes in MG132 treated RPE-1 cells stained for pHSP27, CP110 and PCM1. Six individual z-planes are shown for each example. Scale bars: A, D and E 2 μm; B 10 μm, inset 2 μm. ***p < 0.001.

**Figure S2.**
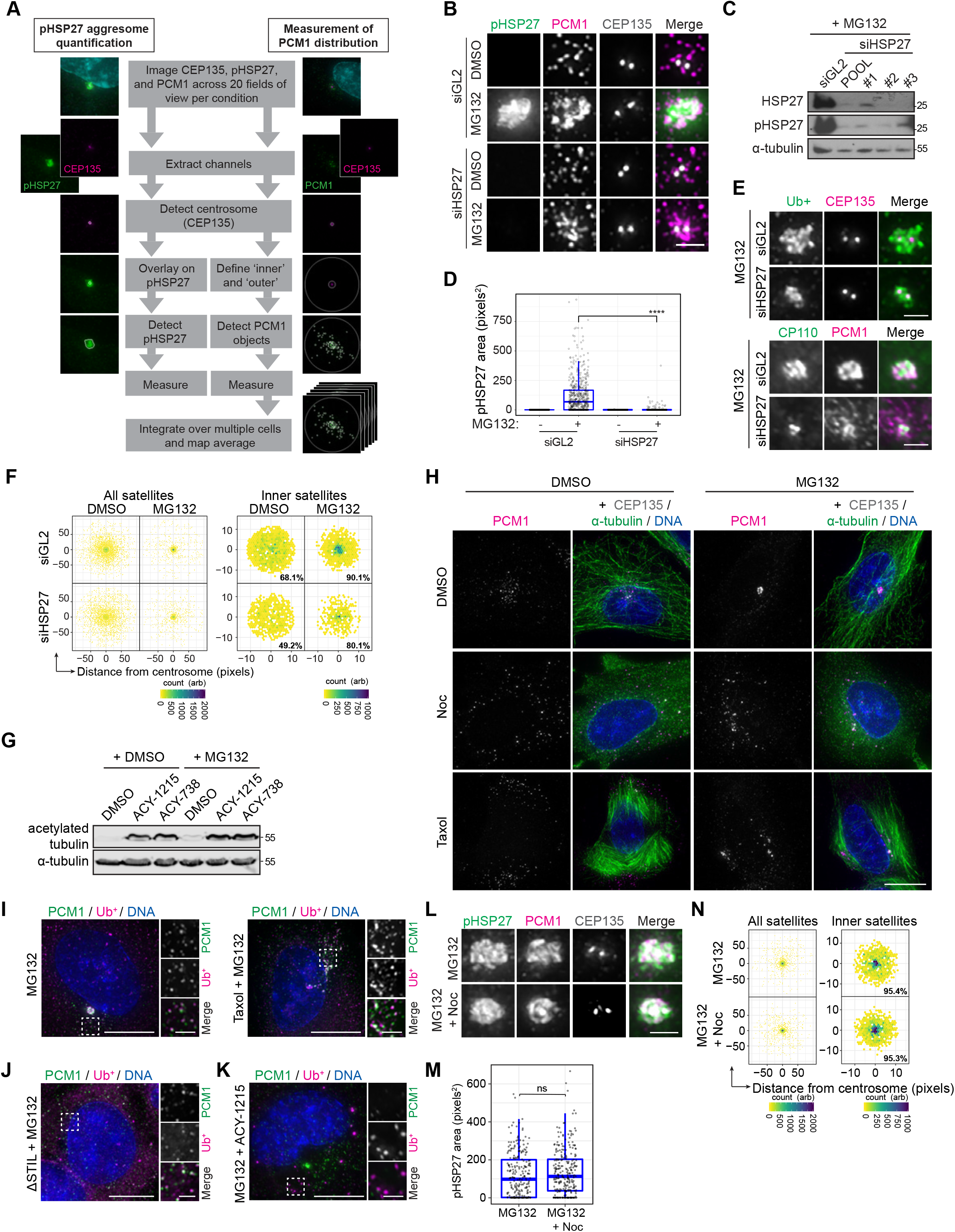
High-resolution quantitative analysis confirms a requirement for protein translation, HDAC6 and microtubules in aggresome formation, related to Figure 2. **A**. Schematic representation of the automated aggresome quantification and satellite mapping pipeline utilized in this study. Cells were stained for pHSP27, PCM1 and CEP135, then at least 20 1536 x 1024 fields of view were captured with 30 z-steps of 0.2 μm per repeat for each condition. Following deconvolution and maximum intensity projection, the images were analyzed using CellProfiler. For aggresome quantification, the channels were extracted then the centrosome detected using the CEP135 channel, the centrosome region was then overlaid on the pHSP27 channel and pHSP27 signal in that area detected and measured. For satellite mapping, following extraction of the channels, an ‘inner’ and ‘outer’ region was defined around the detected CEP135 signal that had been overlaid on the PCM1 channel, then PCM1 objects in these regions detected and measured. **B**. Control (GL2) or HSP27 depleted RPE-1 cells were treated with DMSO or MG132 and stained with antibodies against pHSP27, PCM1 and CEP135. **C**. Immunoblot of cell extracts from control (GL2) and HSP27 knockdown cells treated with MG132 and probed with antibodies against HSP27 and pHSP27. α-tubulin was used as a loading control. **D**. Box-and- whisker plot showing the area occupied by pHSP27 in cells treated as indicated. **E**. Control (GL2) or HSP27 depleted cells treated with MG132 were stained with the indicated antibodies. **F**. Intensity maps of PCM1 distribution relative to the centrosome in cells treated as indicated. The percentage PCM1 signal residing in the defined pericentrosomal region is indicated on the ‘inner’ plots. **G**. Immunoblot of extracts from cells treated with MG132 and HDAC6 inhibitors as indicated, and probed with antibodies against acetylated tubulin and α-tubulin. **H**. RPE-1 cells treated with nocodazole (noc) and taxol +/− MG132 were fixed and stained for PCM1, CEP135 and α-tubulin. DNA was stained with DAPI. **I**. RPE-1 cells treated with MG132 +/− taxol were stained for PCM1 and Ub^+^ proteins. DNA was stained with DAPI. **J**. STIL KO cells were treated with MG132 for 5 hours, then stained for PCM1 and Ub^+^ proteins. DNA was stained with DAPI. **K**. RPE-1 cells treated with MG132 and the HDAC6 inhibitor ACY-1215 were stained for Ub^+^ proteins and PCM1. DNA was stained with DAPI. **L**. RPE-1 cells treated with MG132 or MG132 then nocodazole were stained for pHSP27, PCM1 and CEP135. **M**. Box-and-whisker plot showing the area occupied by pHSP27 in cells treated as in L. **N**. Intensity maps of PCM1 distribution relative to the centrosome in cells treated as indicated. The percentage PCM1 signal residing in the defined pericentrosomal region is indicated on the ‘inner’ plots. Scale bars: B, E and L 2 μm; H, I, J and K 10 μm, insets 2 μm. ****p < 0.0001; ns, not significant.

**Figure S3.**
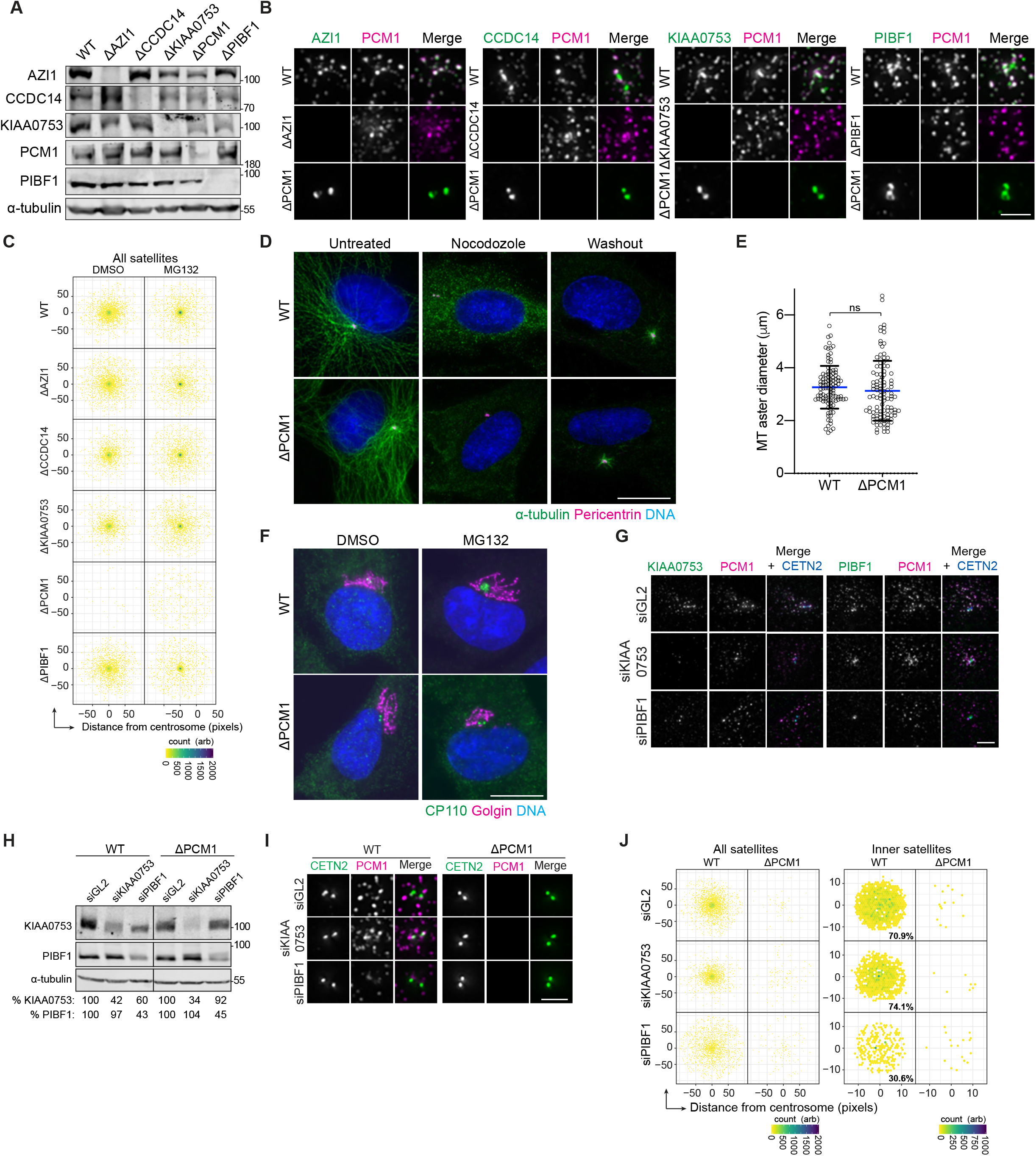
Centriolar satellites are required for aggresome formation, related to Figure 3. **A**. CRISPR/Cas9 mediated gene disruption was used to generate AZI1, CCDC14, KIAA0753, PCM1 and PIBF1 knockout (KO) RPE-1 cell lines. Extracts from WT and KO cells were probed with antibodies against the indicated proteins to confirm loss in the respective cell line. α-tubulin was used as a loading control. **B**. Loss of protein in AZI1, CCDC14, KIAA0753, PCM1 and PIBF1 KO cells was also confirmed by IF microscopy. Untreated cells were stained as indicated. **C**. Intensity maps of PCM1 distribution relative to the centrosome in WT and KO cells treated with DMSO or MG132 for 5 hours. **D**. Staining of α-tubulin, pericentrin and DNA in untreated WT and PCM1 KO cells or those treated with nocodazole for 5 hours with and without 60 seconds washout to allow microtubule regrowth. ns, not significant. **E**. Box-and-whisker plot of the diameter of microtubule (MT) asters formed after 60 seconds of regrowth in WT and PCM1 KO cells. **F**. Staining of CP110, Golgin and DNA in WT and PCM1 KO cells treated with DMSO or MG132. **G**. Staining of KIAA0753 or PIBF1, alongside PCM1 and CETN2 in WT cells treated with siRNAs against GL2 (control), KIAA0753 or PIBF1. **H**. Immunoblot of extracts from WT and PCM1 KO cell lines depleted of GL2 (control), KIAA0753 or PIBF1 by siRNA. α-tubulin was used as a loading control. Numbers at the bottom indicate the mean band intensity expressed as a percentage of signal of the corresponding control for the indicated proteins. **I**. WT and PCM1 KO cells were depleted of GL2 (control), KIAA0753 or PIBF1 and stained for CETN2 and PCM1. **J**. Intensity maps of PCM1 distribution relative to the centrosome in WT and PCM1 KO cells treated and stained as in I. Scale bars: B, G and I 2 μm; D and F 10 μm. Where indicated, DNA was stained with DAPI.

**Figure S4.**
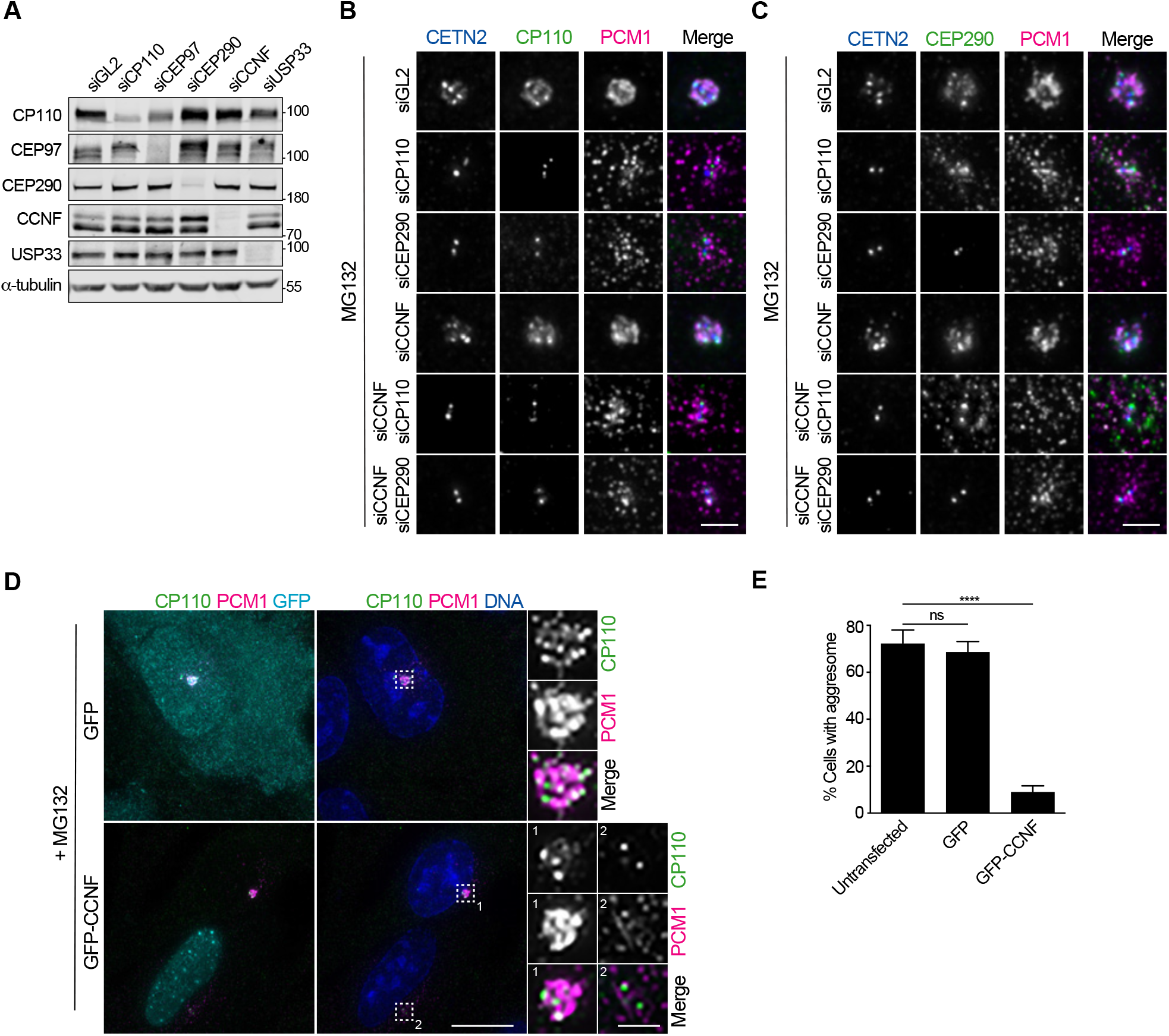
CCNF regulates MG132 induced aggresome formation, related to Figure 4. **A**. Extracts from RPE-1 cells treated with siRNAs against GL2 (control), CP110, CEP97, CEP290, CCNF and USP33 for 48 hours were probed with antibodies to the indicated proteins. α-tubulin was used as a loading control. **B**. RPE-1 cells depleted of GL2, CP110, CEP290, CCNF and CCNF plus CP110 or CCNF plus CEP290 were treated with MG132 and stained for CETN2, CP110 and PCM1. **C**. RPE-1 cells treated as in B were stained for CETN2, CEP290 and PCM1. **D**. RPE-1 cells were transfected with GFP or GFP-CCNF for 24 hours then treated with MG132. Cells were stained for CP110 and PCM1. DNA was stained with DAPI. **E**. Histogram showing the percentage of cells treated as in D that formed an aggresome. Scale bars: B and C 2 μm. D 10 μm, inset 2 μm. ****p < 0.0001; ns, not significant.

**Figure S5.**
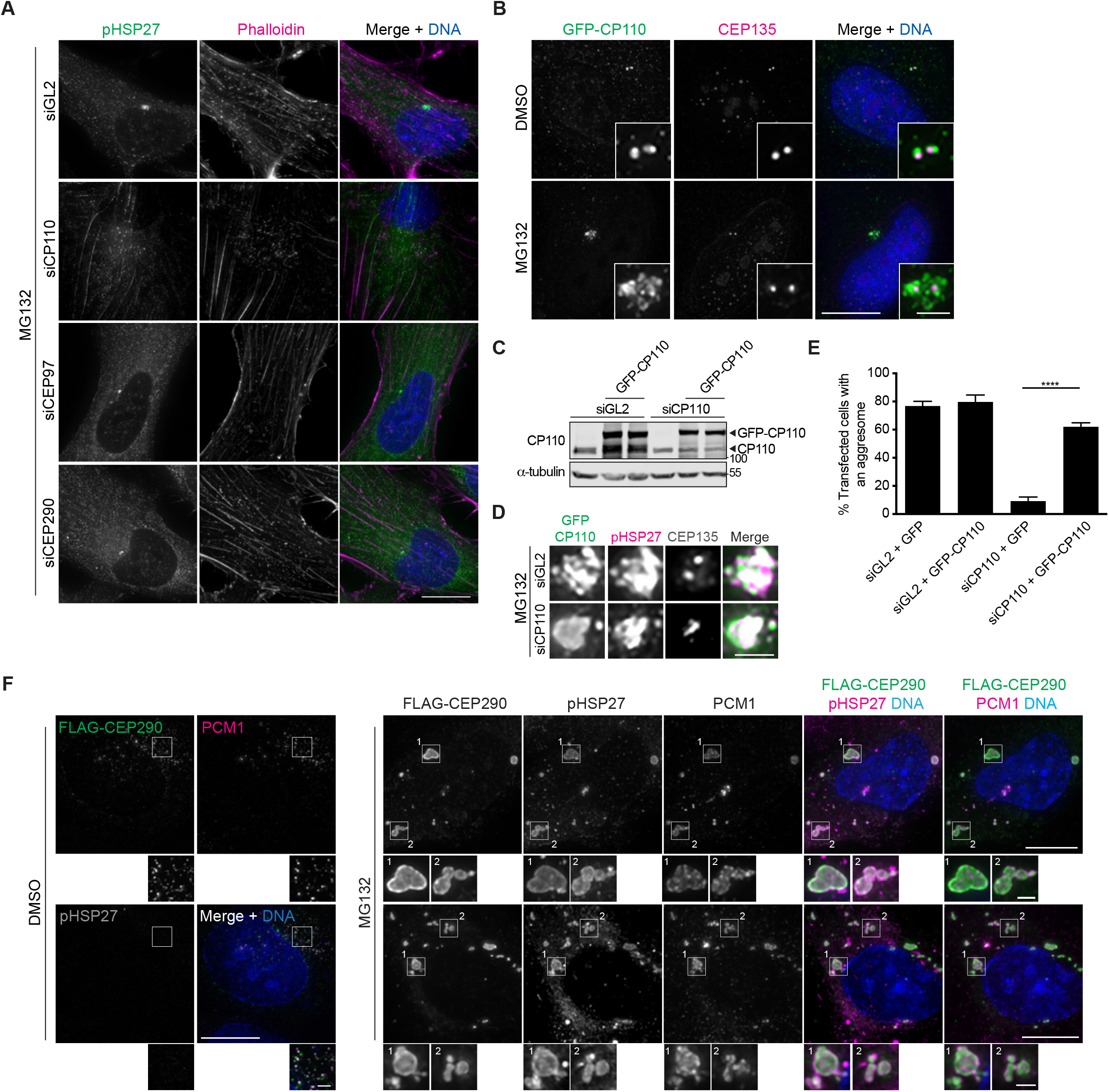
A CP110-CEP97-CEP290 module is required for aggresome formation, related to Figure 5. RPE-1 cells depleted of GL2 (control), CP110, CEP97 or CEP290 for 48 hours were treated with MG132 before being fixed with PFA and stained for pHSP27, actin (phalloidin) and DNA. RPE-1 cells were transiently transfected with GFP-CP110 for 24 hours, then treated with DMSO or MG132 and stained for CEP135 and DNA. **C**. Immunoblot of extracts from WT cells knocked down of GL2 (control) or CP110 for 48 hours and transiently transfected with RNAi-resistant GFP-CP110 for 24 hours. Blots were probed with antibodies against CP110 and α-tubulin as loading control. **D**. siGL2 (control) or siCP110 knockdown cells were transfected with siRNA-resistant GFP-CP110 for 24 hours and MG132 for 5 hours. Cells were stained for pHSP27 and CEP135. **E**. Histogram of the percentage of transfected cells treated as in D that were able to form an aggresome after the addition of MG132. **F**. Cells transfected with FLAG-CEP290 for 24 hours and treated with DMSO or MG132 as indicated for 5 hours were stained for FLAG, PCM1, pHSP27 and DNA. Scale bars: A, B, and F 10 μm, insets B and F 2 μm; D 2 μm. Where indicated, DNA was stained with DAPI. ****p < 0.0001.

**Figure S6.**
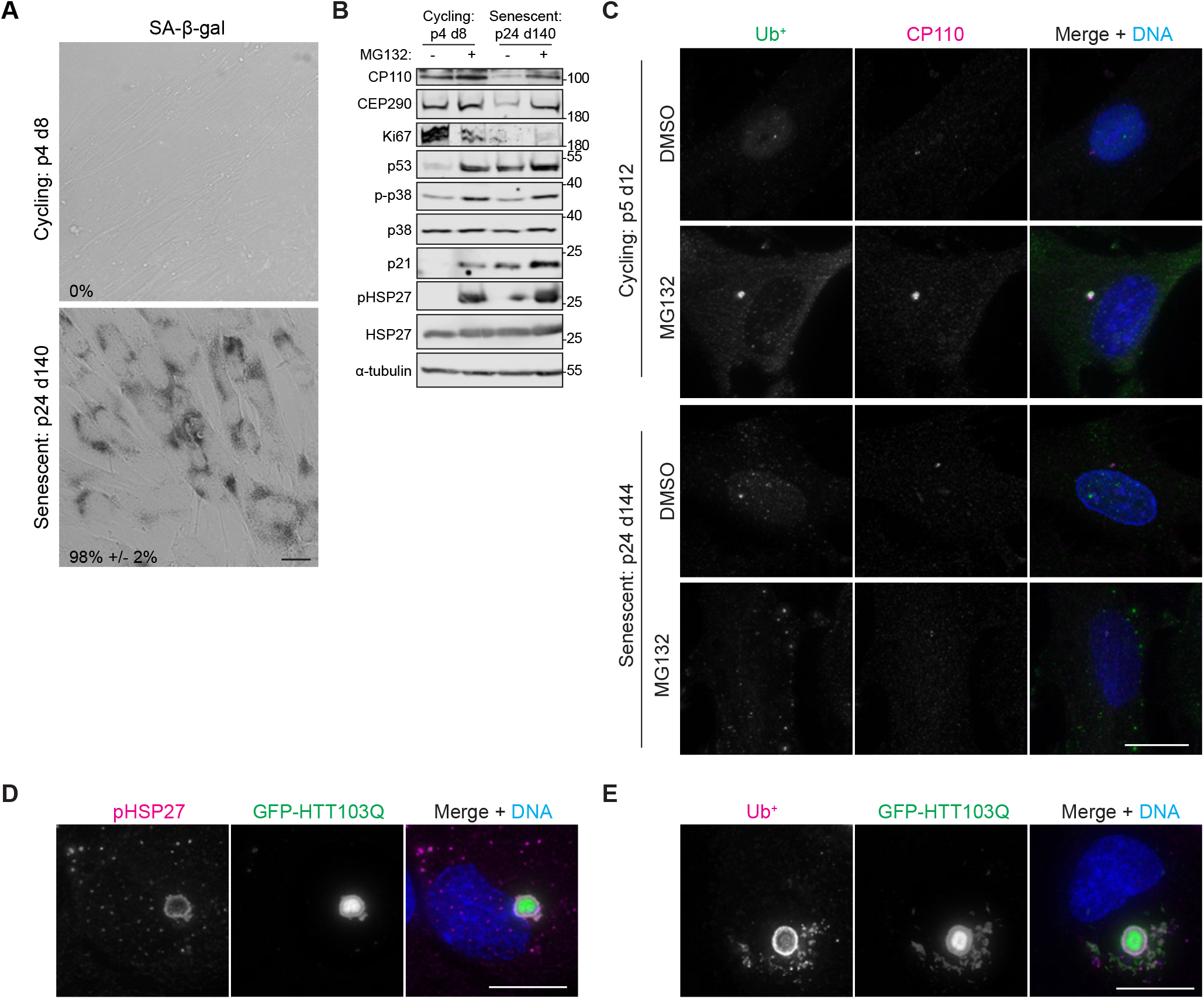
Senescent cells have a reduced capacity to form aggresomes and HTT-polyQ aggregation requires centriolar satellites and the CP110-CEP97-CEP290 module, related to Figure 6. **A**. Bright-field image of cycling and senescent HFF-1 cells that were subjected to a senescence associated β-galactosidase (SA-β-gal) assay. The percentage cells displaying SA-β-gal activity is indicated. **B**. Extracts from cycling and senescent HFF-1 cells treated with or without MG132 were probed for CP110, CEP290, Ki67, p53, phospho-p38, total p38, p21, pHSP27 and total HSP27, as indicated. α-tubulin was used as a loading control. **C**. Cycling and senescent HFF-1 were treated with DMSO or MG132, then stained for Ub^+^ proteins and CP110. DNA was stained with DAPI. **D**. and **E**. RPE-1 cells were transiently transfected with GFP-HTT103Q and stained for pHSP27 (D) or Ub^+^ proteins (E), as indicated. DNA was stained with DAPI. Scale bars: A, C, D and E 10 μm.

**Table S1.**
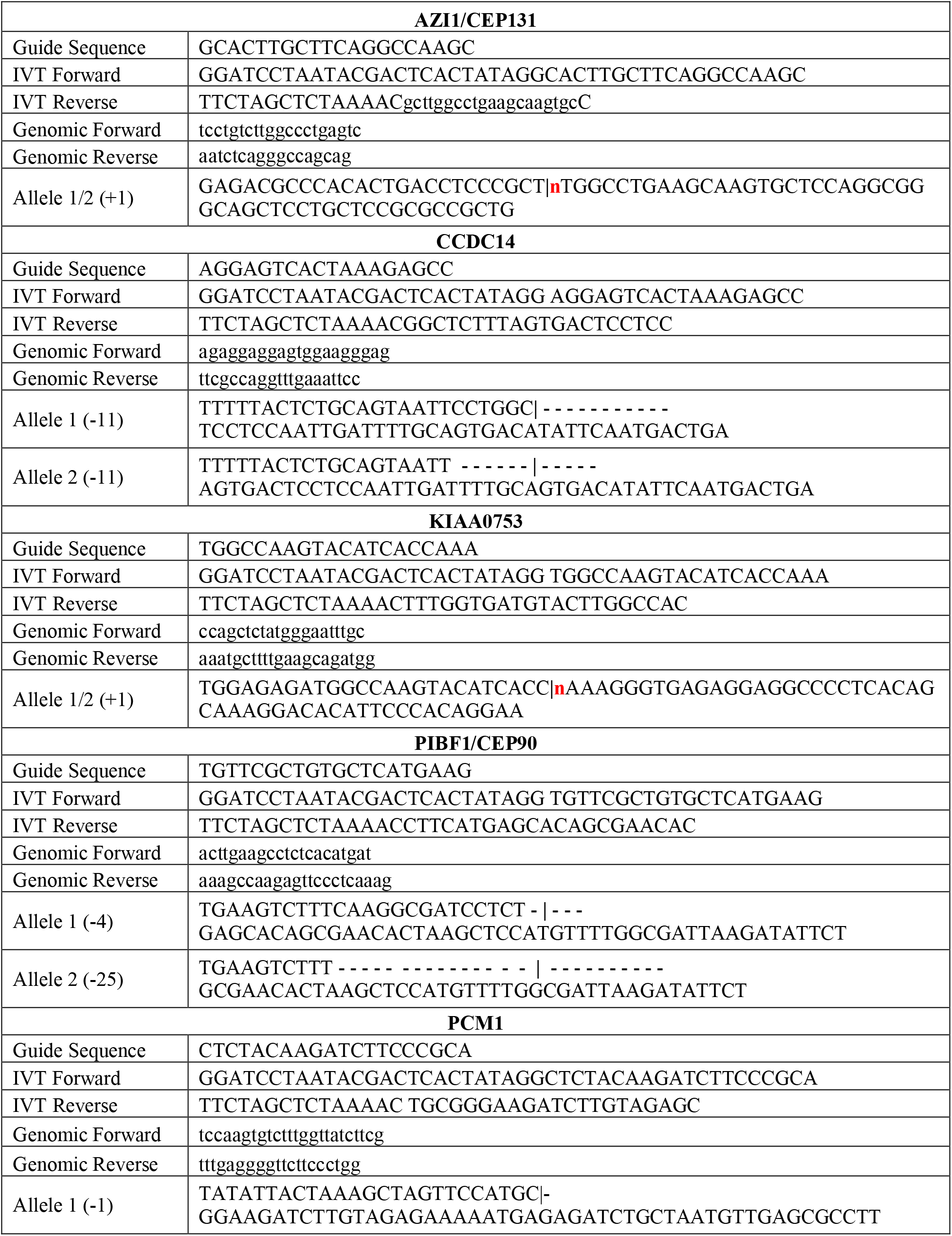
Sequence information for centriolar satellite protein knockout cell lines.

**Table S2.**
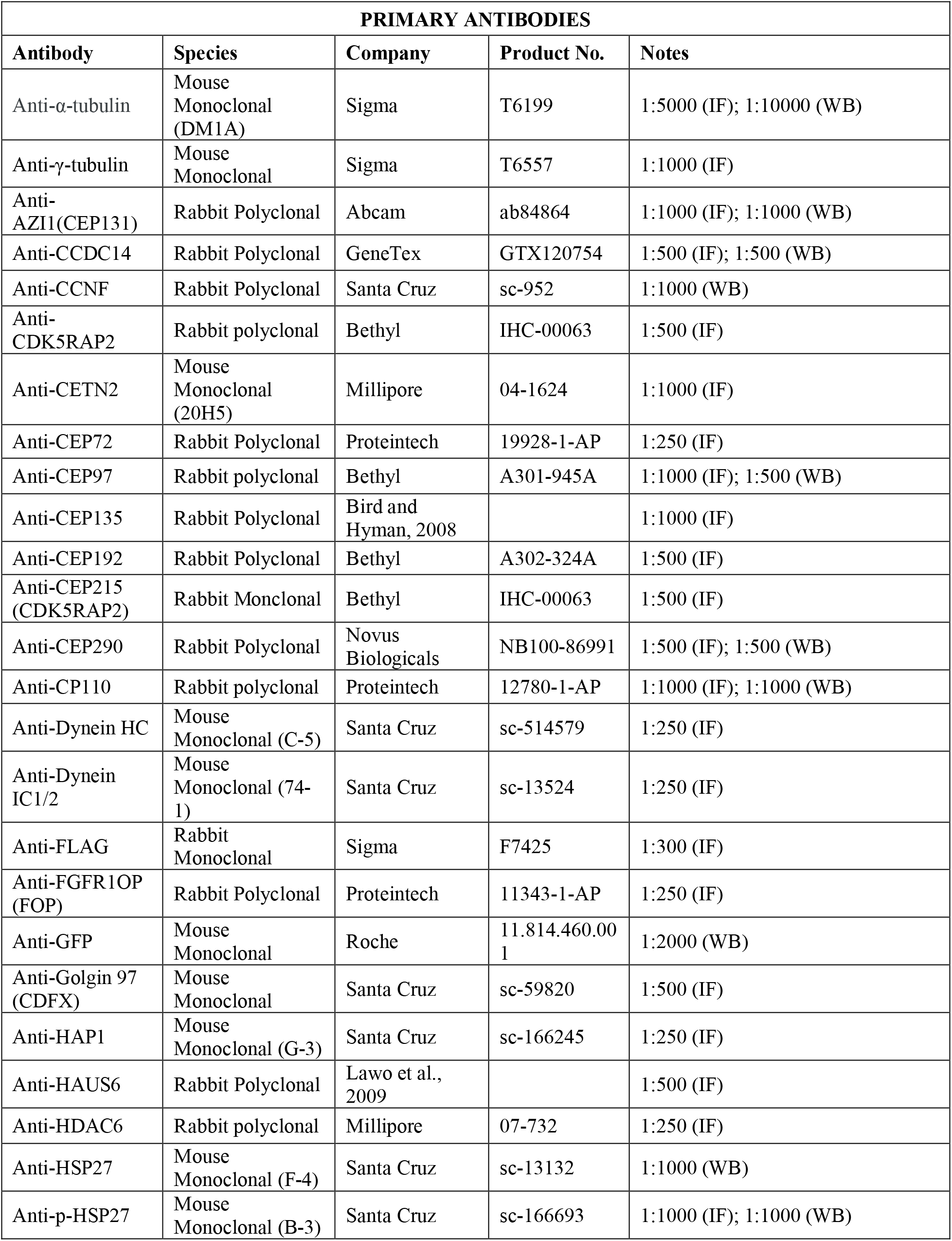
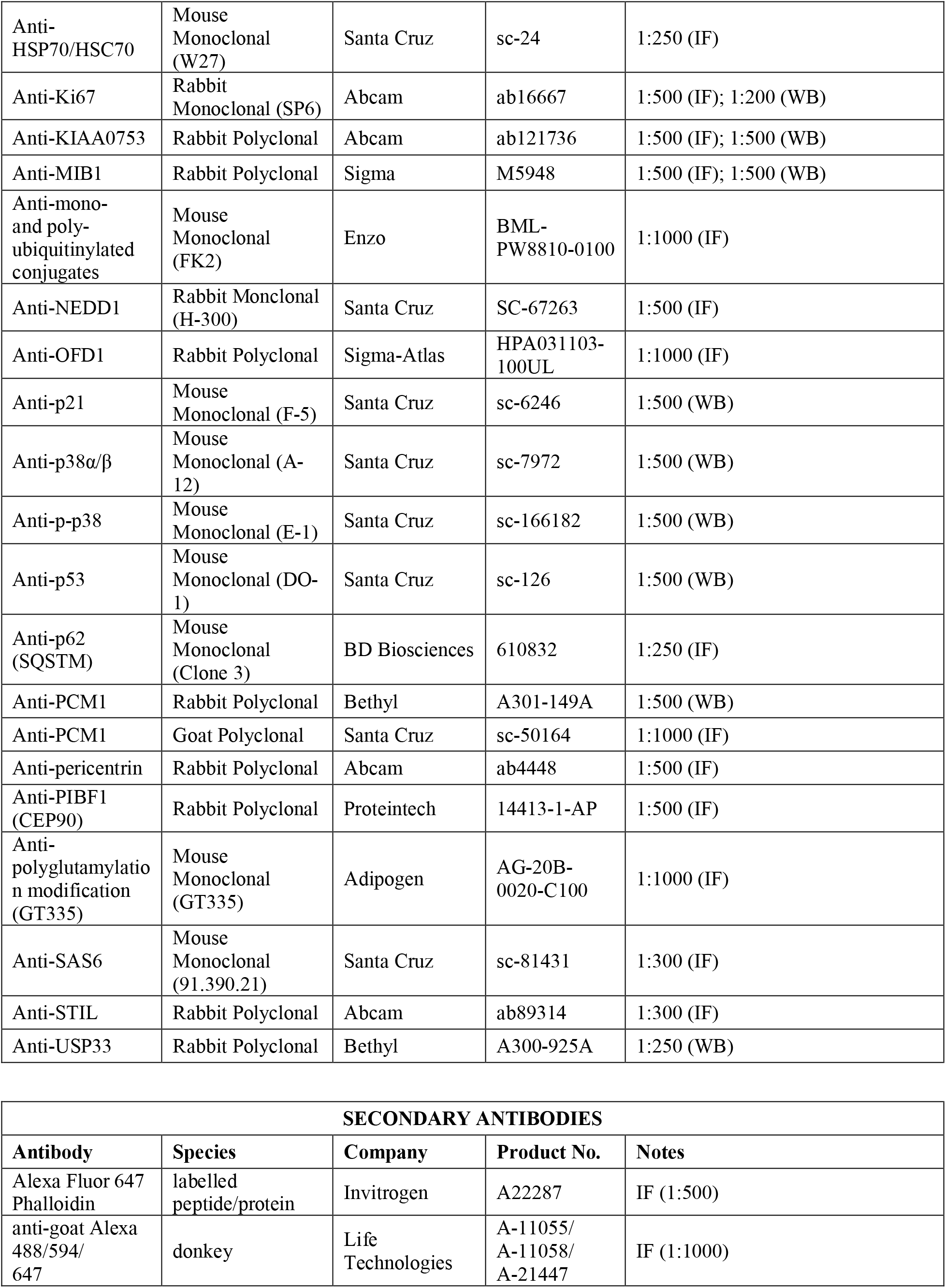
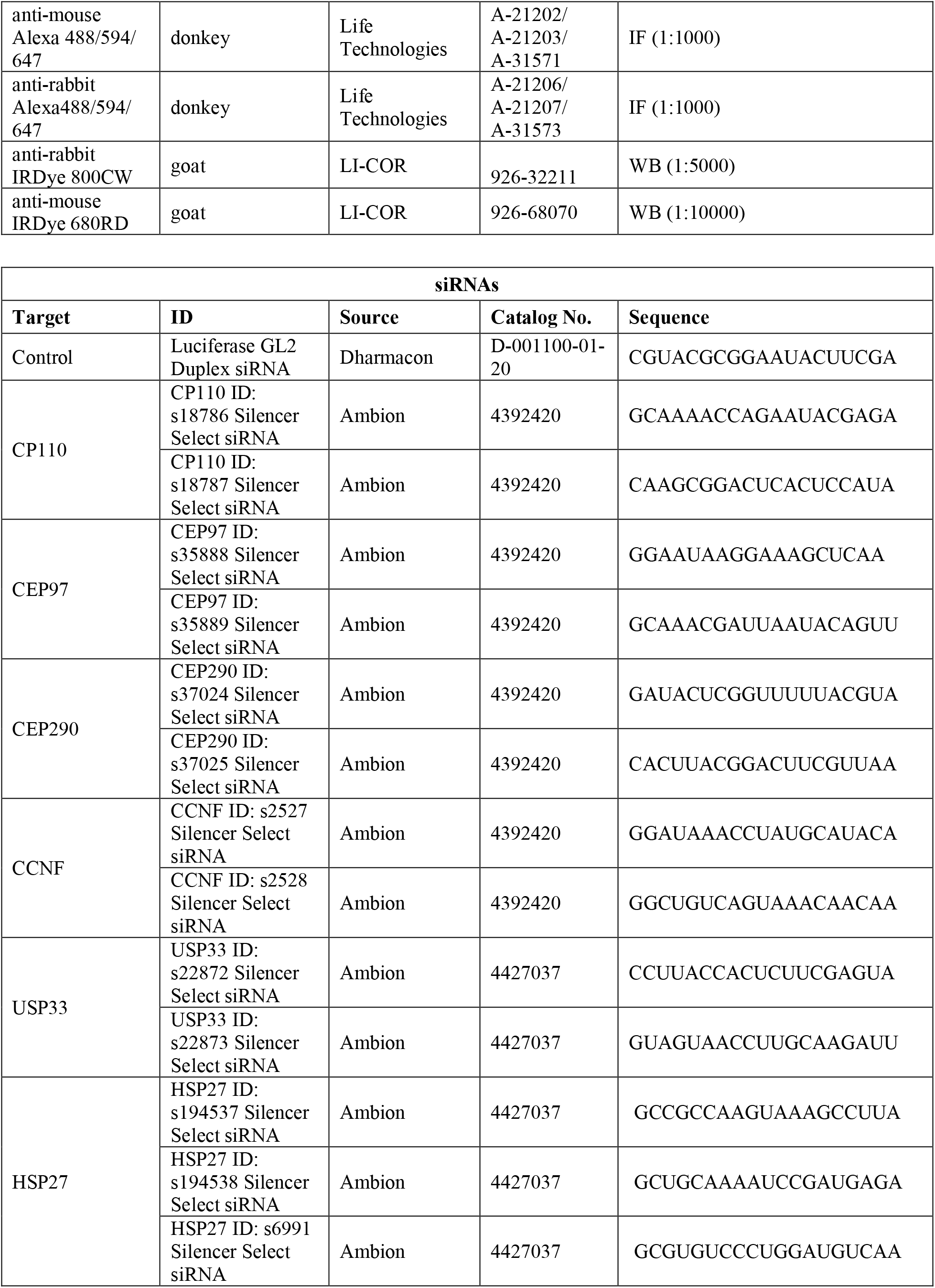
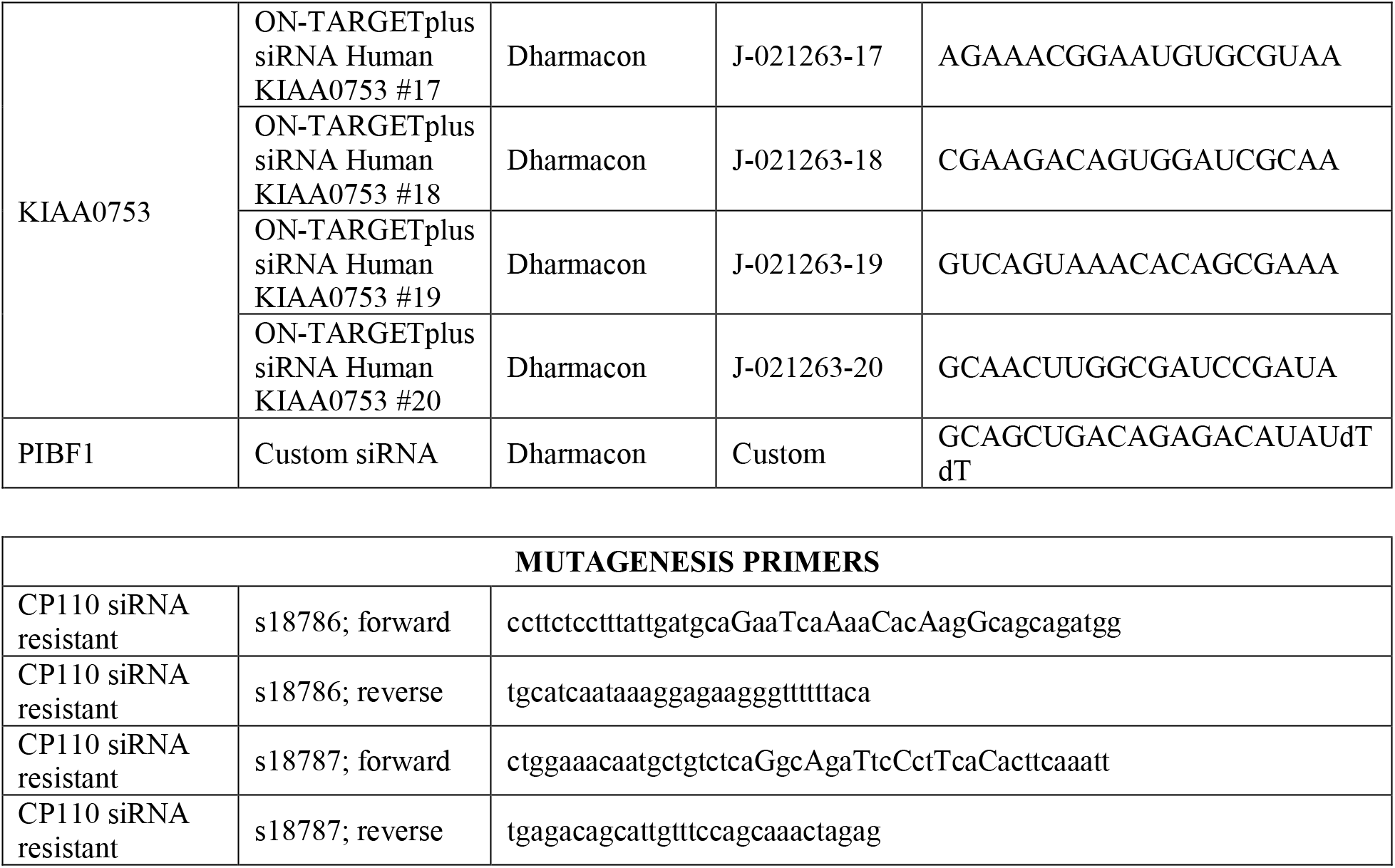
Reagents used in this study.

## REFERENCES

Adegbuyiro, A., Sedighi, F., Pilkington, A.W., IV, Groover, S., and Legleiter, J. (2017). Proteins Containing Expanded Polyglutamine Tracts and Neurodegenerative Disease. Biochemistry 56, 1199–1217.

Antón, L.C., Schubert, U., Bacík, I., Princiotta, M.F., Wearsch, P.A., Gibbs, J., Day, P.M., Realini, C., Rechsteiner, M.C., Bennink, J.R., et al. (1999). Intracellular localization of proteasomal degradation of a viral antigen. J. Cell Biol. 146, 113–124.

Arquint, C., Sonnen, K.F., Stierhof, Y.D., and Nigg, E.A. (2012). Cell-cycle-regulated expression of STIL controls centriole number in human cells. Journal of Cell Science 125, 1342–1352.

Bauer, N.G., and Richter-Landsberg, C. (2006). The dynamic instability of microtubules is required for aggresome formation in oligodendroglial cells after proteolytic stress. J Mol Neurosci 29, 153–168.

Bolhuis, S., and Richter-Landsberg, C. (2010). Effect of proteasome inhibition by MG-132 on HSP27 oligomerization, phosphorylation, and aggresome formation in the OLN-93 oligodendroglia cell line. Journal of Neurochemistry 581, 3665–12.

Breslin, L., Prosser, S.L., Cuffe, S., and Morrison, C.G. (2014). Ciliary abnormalities in senescent human fibroblasts impair proliferative capacity. Cell Cycle 13, 2773–2779.

Chiba, Y., Takei, S., Kawamura, N., Kawaguchi, Y., Sasaki, K., Hasegawa-Ishii, S., Furukawa, A., Hosokawa, M., and Shimada, A. (2012). Immunohistochemical localization of aggresomal proteins in glial cytoplasmic inclusions in multiple system atrophy. Neuropathology and Applied Neurobiology 38, 559–571.

Choi, W.H., Yun, Y., Park, S., Jeon, J.H., Lee, J., Lee, J.H., Yang, S.-A., Kim, N.-K., Jung, C.H., Kwon, Y.T., et al. (2020). Aggresomal sequestration and STUB1-mediated ubiquitylation during mammalian proteaphagy of inhibited proteasomes. Proc. Natl. Acad. Sci. U.S.a. 117, 19190–19200.

Clute, P., and Pines, J. (1999). Temporal and spatial control of cyclin B1 destruction in metaphase. Nat Cell Biol 1, 82–87.

Conduit, P.T., Wainman, A., and Raff, J.W. (2015). Centrosome function and assembly in animal cells. Nat Rev Mol Cell Biol 16, 611–624.

Conkar, D., Culfa, E., Odabasi, E., Rauniyar, N., Yates, J.R., and Firat-Karalar, E.N. (2017). The centriolar satellite protein CCDC66 interacts with CEP290 and functions in cilium formation and trafficking. Journal of Cell Science 130, 1450–1462.

Cuanalo-Contreras, K., Mukherjee, A., and Soto, C. (2013). Role of Protein Misfolding and Proteostasis Deficiency in Protein Misfolding Diseases and Aging. International Journal of Cell Biology 2013, 1–10.

Cunha-Ferreira, I., Rodrigues-Martins, A., Bento, I., Riparbelli, M., Zhang, W., Laue, E., Callaini, G., Glover, D.M., and Bettencourt-Dias, M. (2009). The SCF/Slimb Ubiquitin Ligase Limits Centrosome Amplification through Degradation of SAK/PLK4. Current Biology 19, 43–49.

Čajánek, L., and Nigg, E.A. (2014). Cep164 triggers ciliogenesis by recruiting Tau tubulin kinase 2 to the mother centriole. Proc. Natl. Acad. Sci. U.S.a. 111, E2841–E2850.

Dammermann, A., and Merdes, A. (2002). Assembly of centrosomal proteins and microtubule organization depends on PCM-1. J. Cell Biol. 159, 255–266.

Didier, C., Merdes, A., Gairin, J.-E., and Jabrane-Ferrat, N. (2008). Inhibition of proteasome activity impairs centrosome-dependent microtubule nucleation and organization. Mol. Biol. Cell 19, 1220–1229.

Dimri, G.P., Lee, X., Basile, G., Acosta, M., Scott, G., Roskelley, C., Medrano, E.E., Linskens, M., Rubelj, I., and Pereira-Smith, O. (1995). A biomarker that identifies senescent human cells in culture and in aging skin in vivo. Proc. Natl. Acad. Sci. U.S.a. 92, 9363–9367.

Douanne, T., André-Grégoire, G., Thys, A., Trillet, K., Gavard, J., and Bidère, N. (2019). CYLD Regulates Centriolar Satellites Proteostasis by Counteracting the E3 Ligase MIB1. Cell Reports 27, 1657–1665.e4.

Drummond, D.A. (2012). How Infidelity Creates a Sticky Situation. Molecular Cell 48, 663–664.

Duensing, A., Liu, Y., Perdreau, S.A., Kleylein-Sohn, J., Nigg, E.A., and Duensing, S. (2007). Centriole overduplication through the concurrent formation of multiple daughter centrioles at single maternal templates. Oncogene 26, 6280–6288.

D’Angiolella, V., Donato, V., Vijayakumar, S., Saraf, A., Florens, L., Washburn, M.P., Dynlacht, B., and Pagano, M. (2010). SCFCyclin F controls centrosome homeostasis and mitotic fidelity through CP110 degradation. Nature 466, 138–142.

Engelender, S., Sharp, A.H., Colomer, V., Tokito, M.K., Lanahan, A., Worley, P., Holzbaur, E.L., and Ross, C.A. (1997). Huntingtin-associated protein 1 (HAP1) interacts with the p150Glued subunit of dynactin. Human Molecular Genetics 6, 2205–2212.

Fabunmi, R.P., Wigley, W.C., Thomas, P.J., and DeMartino, G.N. (2000). Activity and regulation of the centrosome-associated proteasome. J. Biol. Chem. 275, 409–413.

Fernández-Cruz, I., and Reynaud, E. (2020). Proteasome Subunits Involved in Neurodegenerative Diseases. Arch Med Res.

Fortun, J., Dunn, W.A., Joy, S., Li, J., and Notterpek, L. (2003). Emerging role for autophagy in the removal of aggresomes in Schwann cells. J Neurosci 23, 10672–10680.

Franz, A., Roque, H., Saurya, S., Dobbelaere, J., and Raff, J.W. (2013). CP110 exhibits novel regulatory activities during centriole assembly in Drosophila. J. Cell Biol. 203, 785–799.

Fuentealba, L.C., Eivers, E., Geissert, D., Taelman, V., and De Robertis, E.M. (2008). Asymmetric mitosis: Unequal segregation of proteins destined for degradation. Proc. Natl. Acad. Sci. U.S.a. 105, 7732–7737.

Fuentealba, L.C., Eivers, E., Ikeda, A., Hurtado, C., Kuroda, H., Pera, E.M., and De Robertis, E.M. (2007). Integrating Patterning Signals: Wnt/GSK3 Regulates the Duration of the BMP/Smad1 Signal. Cell 131, 980–993.

Fujinaga, R., Takeshita, Y., Uozumi, K., Yanai, A., Yoshioka, K., Kokubu, K., and Shinoda, K. (2009). Microtubule-dependent formation of the stigmoid body as a cytoplasmic inclusion distinct from pathological aggresomes. Histochem Cell Biol 132, 305–318.

Fung, E., Richter, C., Yang, H.B., Schäffer, I., Fischer, R., Kessler, B.M., Bassermann, F., and D’Angiolella, V. (2018). FBXL13 directs the proteolysis of CEP192 to regulate centrosome homeostasis and cell migration. EMBO Rep 19, 1358.e1–.e16.

Fusco, C., Micale, L., Egorov, M., Monti, M., D’Addetta, E.V., Augello, B., Cozzolino, F., Calcagnì, A., Fontana, A., Polishchuk, R.S., et al. (2012). The E3-Ubiquitin Ligase TRIM50 Interacts with HDAC6 and p62, and Promotes the Sequestration and Clearance of Ubiquitinated Proteins into the Aggresome. PLoS ONE 7, e40440–13.

Garcia-Mata, R., Gao, Y.-S., and Sztul, E. (2002). Hassles with taking out the garbage: aggravating aggresomes. Traffic 3, 388–396.

García-Mata, R., Bebök, Z., Sorscher, E.J., and Sztul, E.S. (1999). Characterization and dynamics of aggresome formation by a cytosolic GFP-chimera. J. Cell Biol. 146, 1239–1254.

Gerdes, J.M., Liu, Y., Zaghloul, N.A., Leitch, C.C., Lawson, S.S., Kato, M., Beachy, P.A., Beales, P.L., DeMartino, G.N., Fisher, S., et al. (2007). Disruption of the basal body compromises proteasomal function and perturbs intracellular Wnt response. Nat Genet 39, 1350–1360.

Gheiratmand, L., Coyaud, E., Gupta, G.D., Laurent, E.M., Hasegan, M., Prosser, S.L., Gonçalves, J., Raught, B., and Pelletier, L. (2019). Spatial and proteomic profiling reveals centrosome-independent features of centriolar satellites. Embo J 38, e101109.

Goetz, S.C., Liem, K.F., Jr., and Anderson, K.V. (2012). The Spinocerebellar Ataxia-Associated Gene Tau Tubulin Kinase 2 Controls the Initiation of Ciliogenesis. Cell 151, 847–858.

Guderian, G., Westendorf, J., Uldschmid, A., and Nigg, E.A. (2010). Plk4 trans-autophosphorylation regulates centriole number by controlling betaTrCP-mediated degradation. Journal of Cell Science 123, 2163–2169.

Hames, R.S., Crookes, R.E., Straatman, K.R., Merdes, A., Hayes, M.J., Faragher, A.J., and Fry, A.M. (2005). Dynamic recruitment of Nek2 kinase to the centrosome involves microtubules, PCM-1, and localized proteasomal degradation. Mol. Biol. Cell 16, 1711–1724.

Han, K.-J., Wu, Z., Pearson, C.G., Peng, J., Song, K., and Liu, C.-W. (2019). Deubiquitylase USP9X maintains centriolar satellite integrity by stabilizing pericentriolar material 1 protein. Journal of Cell Science 132.

Hao, R., Nanduri, P., Rao, Y., Panichelli, R.S., Ito, A., Yoshida, M., and Yao, T.P. (2013). Proteasomes Activate Aggresome Disassembly and Clearance by Producing Unanchored Ubiquitin Chains. Molecular Cell 51, 819–828.

Hayflick, L. (1965). The limited in vitro lifetime of human diploid cell strains. Experimental Cell Research 37, 614–636.

Hoang-Minh, L.B., Deleyrolle, L.P., Nakamura, N.S., Parker, A.K., Martuscello, R.T., Reynolds, B.A., and Sarkisian, M.R. (2016). PCM1 Depletion Inhibits Glioblastoma Cell Ciliogenesis and Increases Cell Death and Sensitivity to Temozolomide. Transl Oncol 9, 392–402.

Holdgaard, S.G., Cianfanelli, V., Pupo, E., Lambrughi, M., Lubas, M., Nielsen, J.C., Eibes, S., Maiani, E., Harder, L.M., Wesch, N., et al. (2019). Selective autophagy maintains centrosome integrity and accurate mitosis by turnover of centriolar satellites. Nature Communications 10, 4176.

Hori, A., Peddie, C.J., Collinson, L.M., and Toda, T. (2015). Centriolar satellite- and hMsd1/SSX2IP-dependent microtubule anchoring is critical for centriole assembly. Mol. Biol. Cell 26, 2005–2019.

Hou, Y., Dan, X., Babbar, M., Wei, Y., Hasselbalch, S.G., Croteau, D.L., and Bohr, V.A. (2019). Ageing as a risk factor for neurodegenerative disease. Nat Rev Neurol 15, 565–581.

Hung, C.-F., Cheng, W.-F., He, L., Ling, M., Juang, J., Lin, C.-T., and Wu, T.-C. (2003). Enhancing Major Histocompatibility Complex Class I Antigen Presentation by Targeting Antigen to Centrosomes. Cancer Res 63, 2393–2398.

Iqbal, A., Baldrighi, M., Murdoch, J.N., Fleming, A., and Wilkinson, C.J. (2020). Alpha-synuclein aggresomes inhibit ciliogenesis and multiple functions of the centrosome. Biol Open 9, bio054338–10.

Joachim, J., Razi, M., Judith, D., Wirth, M., Calamita, E., Encheva, V., Dynlacht, B.D., Snijders, A.P., O’Reilly, N., Jefferies, H.B.J., et al. (2017). Centriolar Satellites Control GABARAP Ubiquitination and GABARAP-Mediated Autophagy. Curr. Biol. 27, 2123–2136.e2127.

Johnston, J.A., Ward, C.L., and Kopito, R.R. (1998). Aggresomes: a cellular response to misfolded proteins. J. Cell Biol. 143, 1883–1898.

Johnston, J.A., Illing, M.E., and Kopito, R.R. (2002). Cytoplasmic dynein/dynactin mediates the assembly of aggresomes. Cell Motil. Cytoskeleton 53, 26–38.

Joukov, V., and De Nicolo, A. (2019). The Centrosome and the Primary Cilium: The Yin and Yang of a Hybrid Organelle. Cells 8.

Kaliszewski, M., Knott, A.B., and Bossy-Wetzel, E. (2015). Primary cilia and autophagic dysfunction in Huntington’s disease. Cell Death Differ 22, 1413–1424.

Kawaguchi, Y., Kovacs, J.J., McLaurin, A., Vance, J.M., Ito, A., and Yao, T.P. (2003). The deacetylase HDAC6 regulates aggresome formation and cell viability in response to misfolded protein stress. Cell 115, 727–738.

Keller, J.N., Hanni, K.B., and Markesbery, W.R. (2000). Possible involvement of proteasome inhibition in aging: implications for oxidative stress. Mech Ageing Dev 113, 61–70.

Keryer, G., Pineda, J.R., Liot, G., Kim, J., Dietrich, P., Benstaali, C., Smith, K., Cordelières, F.P., Spassky, N., Ferrante, R.J., et al. (2011). Ciliogenesis is regulated by a huntingtin-HAP1-PCM1 pathway and is altered in Huntington disease. J. Clin. Invest. 121, 4372–4382.

Kim, A.H., Puram, S.V., Bilimoria, P.M., Ikeuchi, Y., Keough, S., Wong, M., Rowitch, D., and Bonni, A. (2009). A Centrosomal Cdc20-APC Pathway Controls Dendrite Morphogenesis in Postmitotic Neurons. Cell 136, 322–336.

Kim, J., Krishnaswami, S.R., and Gleeson, J.G. (2008). CEP290 interacts with the centriolar satellite component PCM-1 and is required for Rab8 localization to the primary cilium. Human Molecular Genetics 17, 3796–3805.

Kim, J.C., Badano, J.L., Sibold, S., Esmail, M.A., Hill, J., Hoskins, B.E., Leitch, C.C., Venner, K., Ansley, S.J., Ross, A.J., et al. (2004). The Bardet-Biedl protein BBS4 targets cargo to the pericentriolar region and is required for microtubule anchoring and cell cycle progression. Nat Genet 36, 462–470.

Kimura, H., Miki, Y., and Nakanishi, A. (2014). Centrosomes at M phase act as a scaffold for the accumulation of intracellular ubiquitinated proteins. Cell Cycle 13, 1928–1937.

Kleylein-Sohn, J., Westendorf, J., Le Clech, M., Habedanck, R., Stierhof, Y.-D., and Nigg, E.A. (2007). Plk4-Induced Centriole Biogenesis in Human Cells. Dev. Cell 13, 190–202.

Kocaturk, N.M., and Gozuacik, D. (2018). Crosstalk Between Mammalian Autophagy and the Ubiquitin-Proteasome System. Front Cell Dev Biol 6, 712–727.

Kohlmaier, G., Loncarek, J., Meng, X., McEwen, B.F., Mogensen, M.M., Spektor, A., Dynlacht, B.D., Khodjakov, A., and Gönczy, P. (2009). Overly Long Centrioles and Defective Cell Division upon Excess of the SAS-4-Related Protein CPAP. Current Biology 19, 1012–1018.

Korzeniewski, N., Cuevas, R., Duensing, A., and Duensing, S. (2010). Daughter centriole elongation is controlled by proteolysis. Mol. Biol. Cell 21, 3942–3951.

Korzeniewski, N., Zheng, L., Cuevas, R., Parry, J., Chatterjee, P., Anderton, B., Duensing, A., Münger, K., and Duensing, S. (2009). Cullin 1 functions as a centrosomal suppressor of centriole multiplication by regulating polo-like kinase 4 protein levels. Cancer Res 69, 6668–6675.

Kubo, A., Sasaki, H., Yuba-Kubo, A., Tsukita, S., and Shiina, N. (1999). Centriolar satellites: molecular characterization, ATP-dependent movement toward centrioles and possible involvement in ciliogenesis. J. Cell Biol. 147, 969–980.

Lacaille, V.G., and Androlewicz, M.J. (2000). Targeting of HIV-1 Nef to the centrosome: implications for antigen processing. Traffic 1, 884–891.

Lee, J.H., and Gleeson, J.G. (2010). The role of primary cilia in neuronal function. Neurobiology of Disease 38, 167–172.

Li, J., D’Angiolella, V., Seeley, E.S., Kim, S., Kobayashi, T., Fu, W., Campos, E.I., Pagano, M., and Dynlacht, B.D. (2013). USP33 regulates centrosome biogenesis via deubiquitination of the centriolar protein CP110. Nature 495, 255–259.

Li, S.H., Hosseini, S.H., Gutekunst, C.A., Hersch, S.M., Ferrante, R.J., and Li, X.J. (1998). A human HAP1 homologue. Cloning, expression, and interaction with huntingtin. J. Biol. Chem. 273, 19220–19227.

Li, X., Song, N., Liu, L., Liu, X., Ding, X., Song, X., Yang, S., Shan, L., Zhou, X., Su, D., et al. (2017). USP9X regulates centrosome duplication and promotes breast carcinogenesis. Nature Communications 8, 14866.

Liu, W.J., Ye, L., Huang, W.F., Guo, L.J., Xu, Z.G., Wu, H.L., Yang, C., and Liu, H.F. (2016). p62 links the autophagy pathway and the ubiqutin–proteasome system upon ubiquitinated protein degradation. Cellular & Molecular Biology Letters 1–14.

Liu, Y.P., Tsai, I.-C., Morleo, M., Oh, E.C., Leitch, C.C., Massa, F., Lee, B.-H., Parker, D.S., Finley, D., Zaghloul, N.A., et al. (2014). Ciliopathy proteins regulate paracrine signaling by modulating proteasomal degradation of mediators. J. Clin. Invest. 124, 2059–2070.

López-Otín, C., Blasco, M.A., Partridge, L., Serrano, M., and Kroemer, G. (2013). The Hallmarks of Aging. Cell 153, 1194–1217.

Mao, J., Xia, Q., Liu, C., Ying, Z., Wang, H., and Wang, G. (2017). A critical role of Hrd1 in the regulation of optineurin degradation and aggresome formation. Human Molecular Genetics 26, 1877–1889.

Máthé, E., Kraft, C., Giet, R., Deák, P., Peters, J.-M., and Glover, D.M. (2004). The E2-C Vihar Is Required for the Correct Spatiotemporal Proteolysis of Cyclin B and Itself Undergoes Cyclical Degradation. Current Biology 14, 1723–1733.

McNaught, K.S.P., Shashidharan, P., Perl, D.P., Jenner, P., and Olanow, C.W. (2002). Aggresome-related biogenesis of Lewy bodies. European Journal of Neuroscience 16, 2136–2148.

Meriin, A.B., Zaarur, N., and Sherman, M.Y. (2012). Association of translation factor eEF1A with defective ribosomal products generates a signal for aggresome formation. Journal of Cell Science 125, 2665–2674.

Mishra, A., Godavarthi, S.K., Maheshwari, M., Goswami, A., and Jana, N.R. (2009). The ubiquitin ligase E6-AP is induced and recruited to aggresomes in response to proteasome inhibition and may be involved in the ubiquitination of Hsp70-bound misfolded proteins. J. Biol. Chem. 284, 10537–10545.

Nawrocki, S.T., Carew, J.S., Dunner, K., Jr., Boise, L.H., Chiao, P.J., Huang, P., Abbruzzese, J.L., and McConkey, D.J. (2005). Bortezomib Inhibits PKR-Like Endoplasmic Reticulum (ER) Kinase and Induces Apoptosis via ER Stress in Human Pancreatic Cancer Cells. Cancer Res 65, 11510–11519.

Nigg, E.A., and Holland, A.J. (2018). Once and only once: mechanisms of centriole duplication and their deregulation in disease. Nat Rev Mol Cell Biol 19, 297–312.

Odabasi, E., Gul, S., Kavakli, I.H., and Firat-Karalar, E.N. (2019). Centriolar satellites are required for efficient ciliogenesis and ciliary content regulation. EMBO Rep 20.

Olzmann, J.A., Li, L., and Chin, L.S. (2008). Aggresome formation and neurodegenerative diseases: therapeutic implications. Curr Med Chem 15, 47–60.

Puklowski, A., Homsi, Y., Keller, D., May, M., Chauhan, S., Kossatz, U., Grünwald, V., Kubicka, S., Pich, A., Manns, M.P., et al. (2011). The SCF–FBXW5 E3-ubiquitin ligase is regulated by PLK4 and targets HsSAS-6 to control centrosome duplication. Nat Cell Biol 13, 1004–1009.

Puram, S.V., Kim, A.H., and Bonni, A. (2010). An old dog learns new tricks: a novel function for Cdc20-APC in dendrite morphogenesis in neurons. Cell Cycle 9, 482–485.

Puram, S.V., Kim, A.H., Park, H.-Y., Anckar, J., and Bonni, A. (2013). The Ubiquitin Receptor S5a/Rpn10 Links Centrosomal Proteasomes with Dendrite Development in the Mammalian Brain. Cell Reports 4, 19–30.

Qin, S., Jiang, C., and Gao, J. (2019). Transcriptional factor Nrf2 is essential for aggresome formation during proteasome inhibition. Biom Rep 1–12.

Quarantotti, V., Chen, J.X., Tischer, J., Gonzalez Tejedo, C., Papachristou, E.K., D’Santos, C.S., Kilmartin, J.V., Miller, M.L., and Gergely, F. (2019). Centriolar satellites are acentriolar assemblies of centrosomal proteins. Embo J 38, e101082.

Raff, J.W., Jeffers, K., and Huang, J.-Y. (2002). The roles of Fzy/Cdc20 and Fzr/Cdh1 in regulating the destruction of cyclin B in space and time. J. Cell Biol. 157, 1139–1149.

Rogers, G.C., Rusan, N.M., Roberts, D.M., Peifer, M., and Rogers, S.L. (2009). The SCFSlimb ubiquitin ligase regulates Plk4/Sak levels to block centriole reduplication. J. Cell Biol. 184, 225–239.

Rufini, A., Tucci, P., Celardo, I., and Melino, G. (2019). Senescence and aging: the critical roles of p53. 1–15.

Saez, I., and Vilchez, D. (2014). The Mechanistic Links Between Proteasome Activity, Aging and Age-related Diseases. Curr Genomics 15, 38–51.

Santra, M., Dill, K.A., and de Graff, A.M.R. (2019). Proteostasis collapse is a driver of cell aging and death. Proc. Natl. Acad. Sci. U.S.a. 116, 22173–22178.

Schmidt, T.I., Kleylein-Sohn, J., Westendorf, J., Le Clech, M., Lavoie, S.B., Stierhof, Y.-D., and Nigg, E.A. (2009). Control of Centriole Length by CPAP and CP110. Curr. Biol. 19, 1005–1011.

Selimovic, D., Porzig, B.B.O.W., El-Khattouti, A., Badura, H.E., Ahmad, M., Ghanjati, F., Santourlidis, S., Haikel, Y., and Hassan, M. (2013). Bortezomib/proteasome inhibitor triggers both apoptosis and autophagy-dependent pathways in melanoma cells. Cellular Signalling 25, 308–318.

Spektor, A., Tsang, W.Y., Khoo, D., and Dynlacht, B.D. (2007). Cep97 and CP110 suppress a cilia assembly program. Cell 130, 678–690.

Stowe, T.R., Wilkinson, C.J., Iqbal, A., and Stearns, T. (2012). The centriolar satellite proteins Cep72 and Cep290 interact and are required for recruitment of BBS proteins to the cilium. Mol. Biol. Cell 23, 3322–3335.

Strnad, P., Leidel, S., Vinogradova, T., Euteneuer, U., Khodjakov, A., and Gönczy, P. (2007). Regulated HsSAS-6 Levels Ensure Formation of a Single Procentriole per Centriole during the Centrosome Duplication Cycle. Dev. Cell 13, 203–213.

Szebenyi, G., Wigley, W.C., Hall, B., Didier, A., Yu, M., Thomas, P., and Krämer, H. (2007). Hook2 contributes to aggresome formation. BMC Cell Biol. 8, 19–11.

Tsang, W.Y., Bossard, C., Khanna, H., Peränen, J., Swaroop, A., Malhotra, V., and Dynlacht, B.D. (2008). CP110 Suppresses Primary Cilia Formation through Its Interaction with CEP290, a Protein Deficient in Human Ciliary Disease. Dev. Cell 15, 187–197.

van Deursen, J.M. (2014). The role of senescent cells in ageing. Nature 509, 439–446.

Vertii, A., Zimmerman, W., Ivshina, M., and Doxsey, S. (2015). Centrosome-intrinsic mechanisms modulate centrosome integrity during fever. Mol. Biol. Cell 26, 3451–3463.

Vidair, C.A., Doxsey, S.J., and Dewey, W.C. (1993). Heat shock alters centrosome organization leading to mitotic dysfunction and cell death. J. Cell. Physiol. 154, 443–455.

Vora, S., and Phillips, B.T. (2015). Centrosome-Associated Degradation Limits β-Catenin Inheritance by Daughter Cells after Asymmetric Division. Current Biology 25, 1005–1016.

Vora, S.M., and Phillips, B.T. (2016). The benefits of local depletion: The centrosome as a scaffold for ubiquitin-proteasome-mediated degradation. Cell Cycle 15, 2124–2134.

Wang, L., Lee, K., Malonis, R., Sanchez, I., and Dynlacht, B.D. (2016). Tethering of an E3 ligase by PCM1 regulates the abundance of centrosomal KIAA0586/Talpid3 and promotes ciliogenesis. eLife Sciences 5, e12950.

Wang, Q., Tang, Y., Xu, Y., Xu, S., Jiang, Y., Dong, Q., Zhou, Y., and Ge, W. (2017). The X-linked deubiquitinase USP9X is an integral component of centrosome. J. Biol. Chem. 292, 12874–12884.

Wang, X.J., Yu, J., Wong, S.H., Cheng, A.S., Chan, F.K., Ng, S.S., Cho, C.H., Sung, J.J., and Wu, W.K. (2014). A novel crosstalk between two major protein degradation systems. Autophagy 9, 1500–1508.

Wang, X.-X., Wan, R.-Z., and Liu, Z.-P. (2018). Recent advances in the discovery of potent and selective HDAC6 inhibitors. Eur J Med Chem 143, 1406–1418.

Wigley, W.C., Fabunmi, R.P., Lee, M.G., Marino, C.R., Muallem, S., DeMartino, G.N., and Thomas, P.J. (1999). Dynamic association of proteasomal machinery with the centrosome. J. Cell Biol. 145, 481–490.

Wilkinson, C.J., Carl, M., and Harris, W.A. (2009). Cep70 and Cep131 contribute to ciliogenesis in zebrafish embryos. BMC Cell Biol. 10, 688–14.

Wong, E.S.P., Tan, J.M.M., Soong, W.-E., Hussein, K., Nukina, N., Dawson, V.L., Dawson, T.M., Cuervo, A.M., and Lim, K.-L. (2008). Autophagy-mediated clearance of aggresomes is not a universal phenomenon. Human Molecular Genetics 17, 2570–2582.

Wójcik, C., Schroeter, D., Wilk, S., Lamprecht, J., and Paweletz, N. (1996). Ubiquitin-mediated proteolysis centers in HeLa cells: indication from studies of an inhibitor of the chymotrypsin-like activity of the proteasome. Eur. J. Cell Biol. 71, 311–318.

Wu, Q., Li, B., Le Liu, Sun, S., and Sun, S. (2020). Centrosome dysfunction: a link between senescence and tumor immunity. Signal Transduction and Targeted Therapy 1–9.

Wu, W.K.K., Wu, Y.C., Yu, L., Li, Z.J., Sung, J.J.Y., and Cho, C.H. (2008). Induction of autophagy by proteasome inhibitor is associated with proliferative arrest in colon cancer cells. Biochemical and Biophysical Research Communications 374, 258–263.

Yadav, S.P., Sharma, N.K., Liu, C., Dong, L., Li, T., and Swaroop, A. (2016). Centrosomal protein CP110 controls maturation of the mother centriole during cilia biogenesis. Development 143, 1491–1501.

Zhang, X., and Qian, S.-B. (2011). Chaperone-mediated hierarchical control in targeting misfolded proteins to aggresomes. Mol. Biol. Cell 22, 3277–3288.

Zhou, C., Slaughter, B.D., Unruh, J.R., Guo, F., Yu, Z., Mickey, K., Narkar, A., Ross, R.T., McClain, M., and Li, R. (2014a). Organelle-Based Aggregation and Retention of Damaged Proteins in Asymmetrically Dividing Cells. Cell 159, 530–542.

Zhou, L., Wang, H., Chen, D., Gao, F., Ying, Z., and Wang, G. (2014b). p62/Sequestosome 1 Regulates Aggresome Formation of Pathogenic Ataxin-3 with Expanded Polyglutamine. Ijms 15, 14997–15010.

## References

Bird AW, Hyman AA. Building a spindle of the correct length in human cells requires the interaction between TPX2 and Aurora A. J Cell Biol. 2008;182(2):289–300.

Lawo S, Bashkurov M, Mullin M, Ferreria MG, Kittler R, Habermann B, Tagliaferro A, Poser I, Hutchins JR, Hegemann B, Pinchev D, Buchholz F, Peters JM, Hyman AA, Gingras AC, Pelletier L. HAUS, the 8-subunit human Augmin complex, regulates centrosome and spindle integrity. Curr Biol. 2009;19(10):816–26.

